# Individual-Based Modeling of Microbial Communities Integrating Genetic Mechanisms: A Case Study of LuxS-Mediated Quorum Sensing in Salmonella Typhimurium

**DOI:** 10.1101/2025.03.09.642204

**Authors:** Athanasios D. Balomenos, Anastasia Tampakaki, Elias S. Manolakos

## Abstract

We present a digital twin framework for simulating microbial communities at the single-cell level, integrating genetic mechanisms through individual-based modeling (IBM). This in silico approach enables the study of bacterial populations with bio-sensing capabilities and stochastic virulence expression, facilitating the design of biotechnological applications such as targeted drug delivery. By combining kinetic modeling with IBM, we capture the regulatory interplay between quorum sensing (QS) and virulence, allowing for predictive simulations before costly wet-lab experiments with state-of-the-art technologies like organ-on-chip models.

To demonstrate the power of this approach, we focus on Salmonella Typhimurium, where the LuxS/Autoinducer-2 QS system controls the expression of the type three secretion system (TTSS) encoded by Salmonella pathogenicity island-1 (SPI-1). TTSS-1 effectors are not only key virulence factors but also promising tools for precise intracellular delivery of therapeutic agents. These bacterial-derived effectors function as cell-penetrating effectors (CPEs), autonomously translocating into host cells and overcoming major hurdles in pharmacology by enabling targeted drug delivery.

Our digital twin framework enables the simulation of Salmonella cells engineered to sense their environment and dynamically regulate virulence expression for the controlled secretion of effectors, including potential applications in delivering surrogate drugs to cancerous cells. This work establishes an advanced computational platform for optimizing bacterial therapies and accelerating the development of next-generation biomedical solutions.

## Introduction

Bacterial cells own evolved signaling networks enabling them to sense their micro-environment by producing, exporting and importing small signaling molecules, called autoinducers. Autoinducers can rapidly diffuse across cell populations and accumulate over time. Thus bacteria are apt to collect feedback on the cellular density in their surroundings. This feedback can then be exploited so as to generate decentralized population-wide responses at high cell densities. This microbial phenomenon, known as quorum sensing (QS), has been proved important for several microbiological mechanisms given that it was initially served as bioluminescence regulator [1,2]. In particular, it appears to be a key regulator of several bacterial phenotypes with medical and biotechnological implications, e.g. virulence factor production, biofilm development, persister cells formation and synthesis of antibiotics [3,4,5,6,7,8,9,10]. Many types of QS systems have been described in several bacterial species [11]. For instance, a well-known QS pathway, in which LuxS produced autoinducer-2 (AI-2) mediates signaling, has been found in both Gram-negative and Gram-positive bacteria [12]. The LuxS/AI-2 QS (LuxS-QS) system lies in *Salmonella enterica* serovar Typhimurium (*S.* Typhimurium) [13,14], see Fig 1.

**Fig 1.**
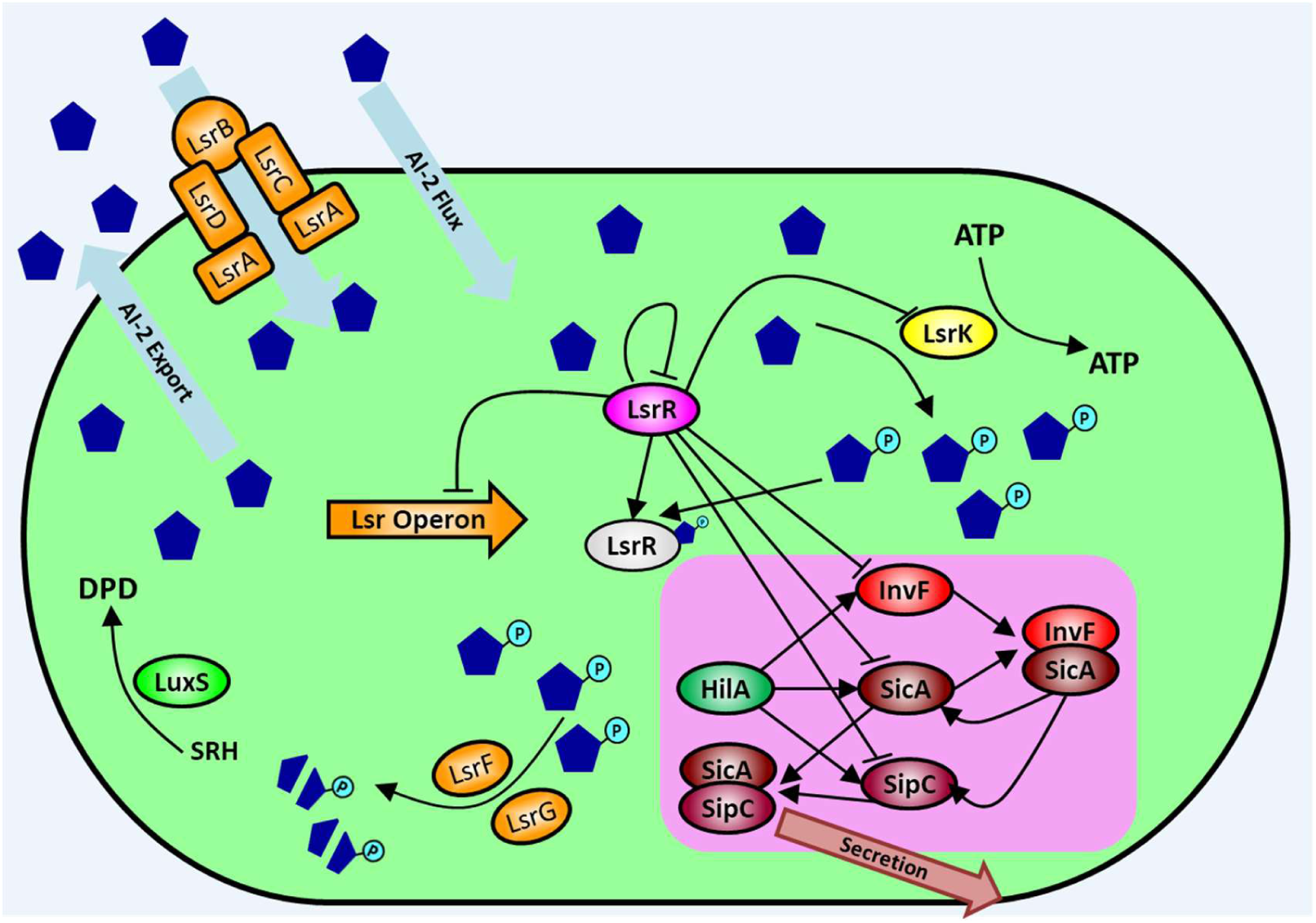
Components and regulatory mechanism of LuxS-QS and TTSS-1 in S. Typhimurium. LsrR initially represses lsr operon (Lsr Operon), lsrRK (LsrR and LsrK) and negatively controls inv operon transcription. As growth ensues extracellular AI-2 (blue pentagons) accumulates in the environment, and cell starts importing it via a PTS-dependent transport (which is not exactly specified [11]) denoted as AI-2 Flux. LsrK phosphorylates AI-2, producing P-AI-2 which begins binding to LsrR. Thus, LsrR stops repressing the lsr genes and the inv operon. Lsr-dependent transport begins; AI-2 binds to the periplasmic binding protein, LsrB, and the Lsr transporter which consists of the two transmembrane proteins, LsrC and LsrD, and the ATP-binding protein LsrA internalizes it (simplified to Lsr Operon species, orange color). More P-AI-2 is formed as the cell densities grow and the transporter’s expression is highly induced. AI-2 internalization increases and extracellular AI-2 is rapidly depleted due to this positive feedback loop. LsrG and LsrF (simplified to Lsr Operon species, orange color) degrade P-AI-2. Simultaneously, if an environmental cue, triggering virulence weaponry (via HilA global regulator), is introduced to the system, on the assumption that needle complex is already formated, InvF is expressed forming a InvF:SicA complex (with basal SicA concentration) which in turn induces the production of SicA (chaperone) and SipC (secreted effector with translocon role). SipC forms a complex with SicA (SicA:SipC) in order to be transferred to the needle complex (not examined in this work) for secretion. Cells reaching the quorum are synchronized to produce and secrete SipC.

*S.* Typhimurium can invade epithelial cells of the small intestine during infection of animal hosts. More specifically, it causes gastroenteritis and diarrhea in humans due to acute intestinal inflammation and a typhoid-like disease in mice. Invasion is accomplished by a type three secretion system which is encoded in Salmonella pathogenicity island 1 (SPI-1). The type three secretion system SPI-1 (TΤSS-1) assembles a needlelike complex via which many effector proteins are translocated into host cells and modify the actin cytoskeleton [15,16]. Several environmental cues which reflect the complex conditions in the intestinal lumen, such as oxygen tension, osmolarity, pH, and nutrients, regulate the expression of SPI-1 [17,18,19], see Fig 1. As proposed in [20] bacterial effectors are very important due to their capability of being exploited as tools for the directed manipulation of host signaling for the host’s benefit. Recently, effector proteins have been identified to autonomously translocate into host cells, representing a novel class of cell-penetrating effectors (CPEs). Moreover, autonomous cell penetration enables transducing specific therapeutic agents to intracellular targets, which is a major hurdle in pharmacology. Another approach that is attracting the attention of both the research community and the pharmaceutical industry includes the use of engineered bacteria as vectors for drug delivery [21,22,23]. As vectors, bacterial cells not only address manufacturing and stability difficulties by synthesizing therapeutics on demand but also may enable targeted delivery. Hopefully, within the next years, their manufacturing and delivery capabilities will be coupled with their natural capacity of bio-sensing and signal integration contributing to the development of both a more intelligent disease monitoring and a controlled dosing process [24].

Generally phenotypic heterogeneity is considered as a dynamic source of diversity in many phenomena [25] such as quorum sensing [26] and virulence [27]. Microbial populations benefit from the creation of variant subpopulations that have the potential to be better equipped in order to persist the environment’s perturbations [25] and exploit new niches [28]. The benefits of heterogeneity to the fitness of the population can be readily envisaged. In vivo experiments to test the reality of such benefits are still lacking, because heterogeneity at the single-cell level is typically masked in conventional studies of microbial populations [25,28], which rely on data averaged across thousands or millions of cells in a sample. However, modelling studies support the hypothesis of heterogeneity’s benefits, at least under certain conditions [29].

Individual cells of microbial pathogens display variable degrees of virulence, have varying degrees of resistance to antimicrobial treatments and other stresses, and exhibit differing propensities to differentiate [25] and to express motility determinants [28]. In addition to the said phenomena, the principal control processes regulating cell function (for example, gene transcription and translation) can, at any moment in time, be differentially activated in different cells of a genetically uniform population. Diversity at the molecular level, is attributed to stochasticity or “noise” in gene transcription and translation [30]. Stochasticity has been understood as a cause of phenotypic heterogeneity [31], typically when heterogeneous phenotypes remain less well defined. Several modeling approaches have been proposed to capture and analyze QS and TTSS-1 dynamics of different bacterial species [32,33,34,35,36]. In [32], Melke *et al.* used an individual-based model of growing bacterial microcolonies to investigate a quorum-sensing mechanism at a single-cell level. They analyzed a molecular network with two positive feedback loops describing the LuxIR circuit in Vibrio fischeri. In [33], Hooshangi and Bentley employed a combination of experimental work and kinetic modeling to decipher network connectivity and signal transduction in the LuxS-QS system of E. coli. In [34], Quan *et al.* used mathematical simulations to examine how desynchronized E. coli’s LuxS-QS activation, arising from cell-to-cell population heterogeneity, could lead to bimodal Lsr signaling and fractional activation. In [35], Saini *et al.* determined the function of S. Typhimurium TTSS-1 regulation circuit, by constructing a kinetic model of the SPI1 gene circuit. In [36], Temme *et al.* built a kinetic model to examine the induction and relaxation dynamics of S. Typhimurium TΤSS-1 mechanism, focusing on the regulation of the effectors’ expression in response to the secretion capacity of the cell. All the aforementioned modelling approaches are deterministic. To the best of our knowledge there is no modelling approach that combines LuxS-QS and TTSS-1 in S. Typhimurium and considers population heterogeneity.

In this work, on the basis of the experimental evidence proposed in [37,38] and the computational modelling we examine the impact of stochasticity on population phenotypes, e.g. virulent, non-virulent, and we recapitulate an interesting biotechnological scenario. First, we propose a kinetic model that combines LuxS-QS with TTSS-1 in *S.* Typhimurium at the population dynamics level (kinetic modeling approach), and then in microbial communities single-cell level (individual-based modeling approach). For this to be done, we formulated a mechanistic mathematical model, see Fig 1, to account for the quorum sensing and virulence mechanism interplay, benefiting from literature knowledge. Using this kinetic model, we manage to recapitulate in silico how quorum sensing mechanism coordinates virulence at population level. We confirmed the mechanistic basis for the LsrR-mediated genetic “coordinator” of S. Typhimurium virulence through QS circuitry, assuming that main virulence weaponry is “on” (due to environment cues).

Then we integrate this model into a multicellular framework following individual-based modeling approach [39,40], to account for noise that exists in single-cell gene expression [41] and growth [42,43] of S. Typhimurium. In the individual-based modeling approach, we exploit the knowledge acquired frοm kinetic model in silico experimentation and we investigate, by perturbating quorum sensing, how virulence is affected in a well-mixed population. To the best of our knowledge in population-based simulations such a system cannot be quantified and reproduced for further analysis and investigation. Cells coordination is buried because it cannot be traced. Noise in growth can introduce noise in virulence expression due to spatiotemporal patterns of AI-2 molecule diffusion. We build an in-silico system, inspired from recent experimental approaches [44,45], in which the species simultaneously produces and transmits AI-2 signal molecules and can become virulent depending on the AI-2 availability in the microenvironment. We show that noise in single-cell growth results in different communication and virulence patterns. Furthermore, we investigate if and to what extent stochasticity in growth can lead into a bistable virulence expression in the population. Finally, we used the proposed framework, as a digital twin, to simulate an interesting biotechnology scenario that can lead to further wet lab experimentation and thus to microbiological breakthroughs. Our modeling results illustrate the importance of noise even in a well-regulated mechanism, such as the LuxS-QS coupled with TTSS-1.

The rest of the paper is organized as follows; In the Results section, we present the devoloped models and their dynamic behaviour. Then, we validate the proposed models and we proceed accordingly to predictions following a constructive presentation of intersting multicellular simulation scenario. In Methods section, we describe the proposed modeling methodology and how we implemented and calibrated each model. On that basis, we elaborate on the personalized growth kinetic parameters used in Ibm approach.

## Results

### Model development

#### Kinetic model LuxS-QS module

First, we formulated a minimal regulatory network to represent LuxS-QS system. As it is claimed in [46], this system is highly dependent on AI-2 concentration, and LsrR repression to all of the network components. So, we used as basis an already perceived network architecture [46,47] to get the most accurate but also parsimonious depiction of the overall QS system behavior. The network contains two negative regulations and a positive feedback loop.

To account for such regulations, we formulated a mathematical model of the Lsr system and its sub-modeules. We model 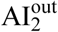, i.e. the AI-2 lying “outside” of the cell membrane, and its downstream regulation. Then, we assume that 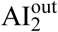 is imported by OP (a transmembrane complex LsrACBD) [46] and PTS-mechanism [46] into the cell. Notice that upon entering the cell, 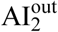 is denoted as AI_2_, which is then phosphorylated by a cytoplasmic kinase, K (LsrK). The 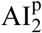 (phosphorylated AI-2) interacts with a transcriptional regulator, R (LsrR) [19]. R inhibits transcription of the OP (lsr-operon) by binding to the lsr-operon promoter site. R also acts as an auto-regulator by binding the lsrR promoter site and inhibiting its own transcription [33]. 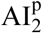 reportedly binds to R protein and prevents inhibition of the lsr-operon, hence alleviating repression and increasing OP production. An increase in the levels of the OP expedites AI_2_ uptake and creates a positive feedback loop where higher concentrations of AI_2_ within the cell result in an increase in AI_2_ uptake. The presence of K is essential for normal operation of this network, thus without this kinase, the system is deactivated, i.e. repressed by R. K phoshorylates AI_2_, producing 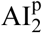 which in turn binds to R and neutralize its action, forming a complex 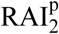 (LsrR:P-AI-2). In Fig 2 (A), we present the reaction network of the QS module. In Table 1, we present the corresponding kinetic laws of the reaction network. The estimated parameters of each kinetic law are presented in S2 Table. The details on the parameter estimation task is provided in Methods Section. R, K and OP were each modeled by a single equation (enclosing protein synthesis, decay, cooperative binding, and repression). 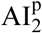 binding to LsrR was modeled as complex formation. This mechanism leads to a system of 6 ordinary differential equations (ODEs) that capture the change in the total number of each protein in the cell. An interested rader can find the system of ODEs and the corresponding rate parameters, in S1 and S3 Table, respectively.

**Fig 2.**
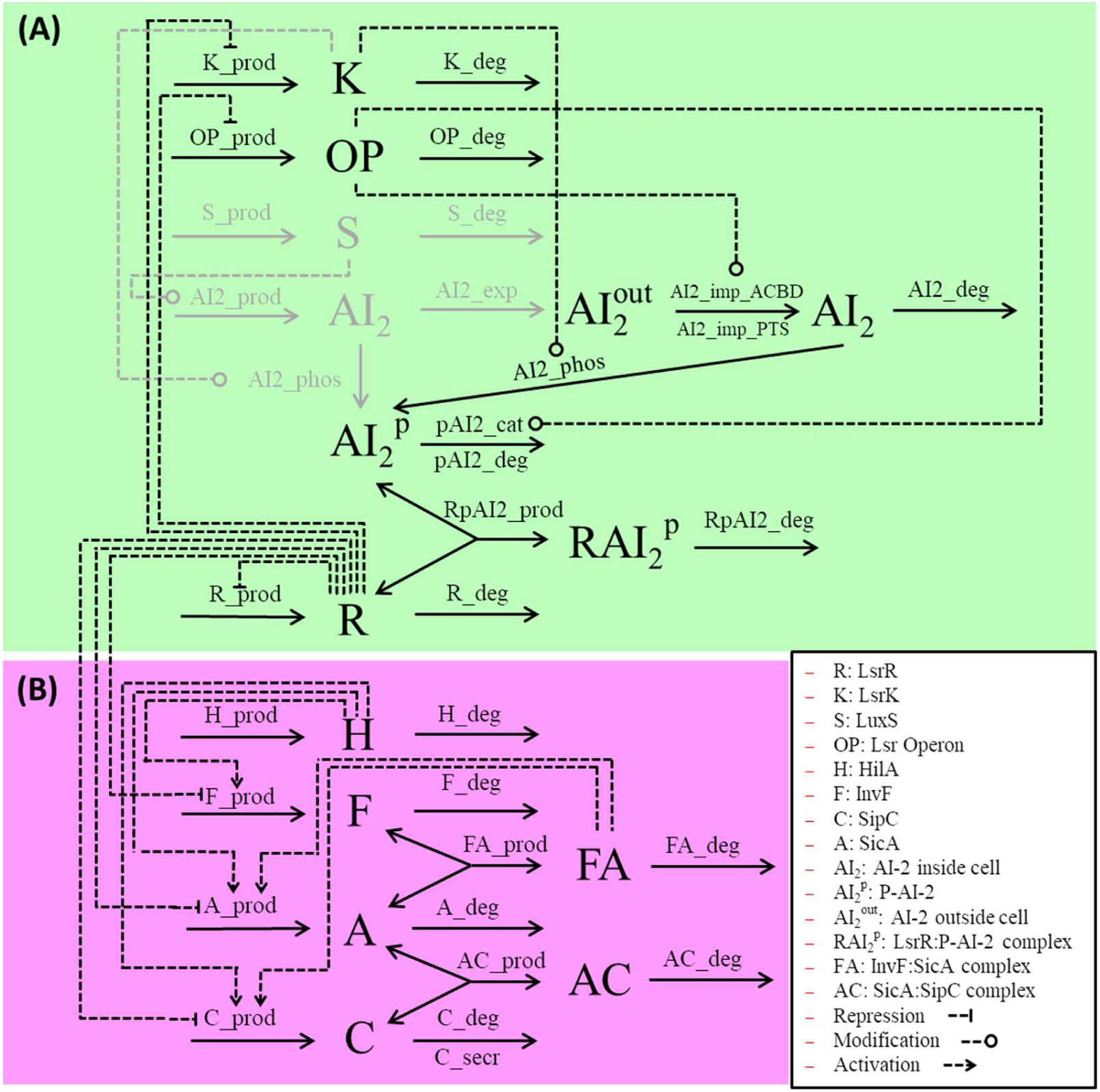
Outline of the reaction network. (A) The kinetic model of LuxS-QS module. (B) The kinetic model of TTSS-1 module. In the legend, we present the abbreviations of species names and the different regulation types found in our model. Above each reaction’s arrow we denote the symbols of the corresponding kinetic laws. See Table 1 for the mathematical expression of each reaction’s kinetic law. Species and reactions in gray participate only in individual based modelling approach (see text for details).

**Table 1.**
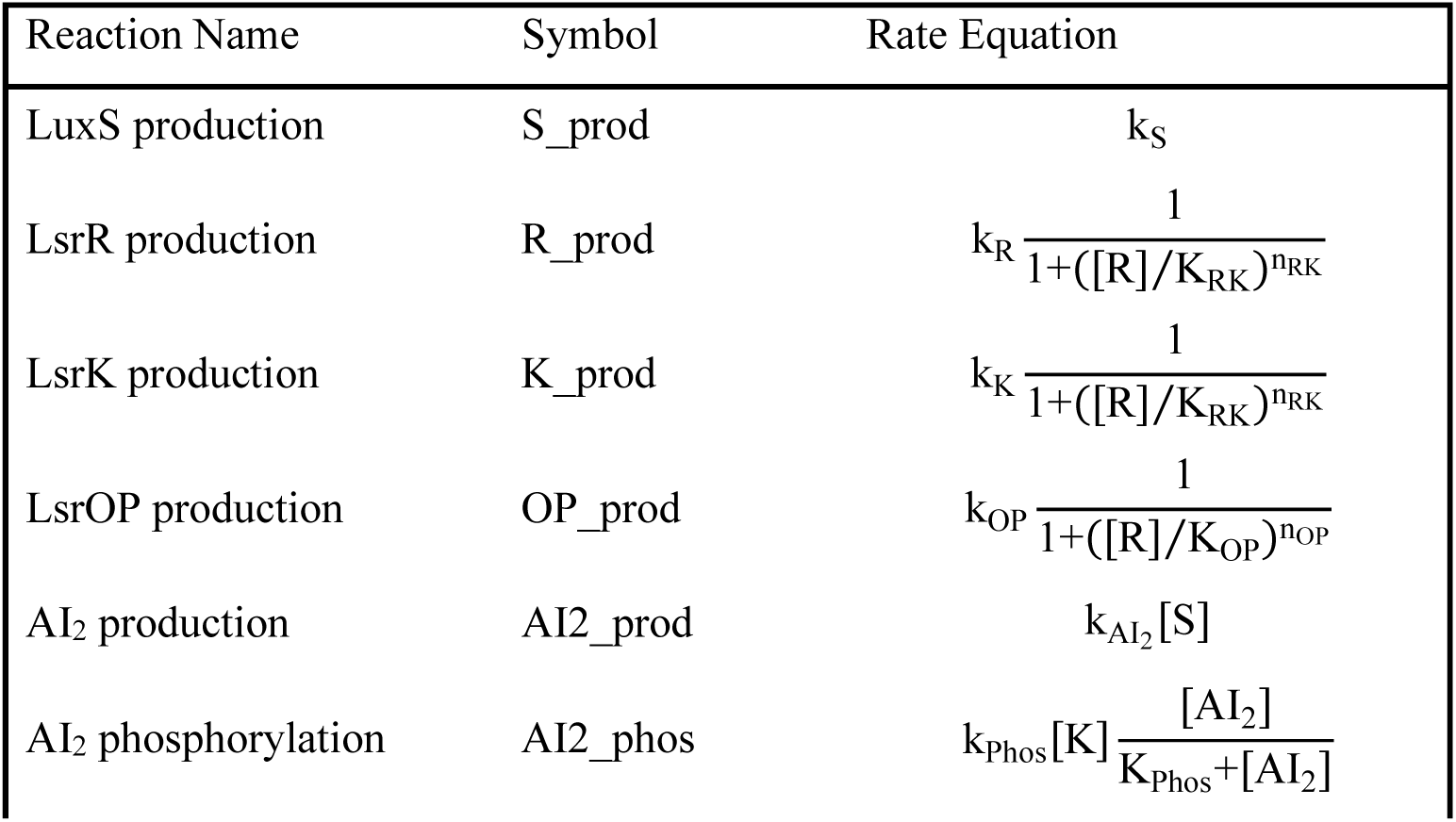
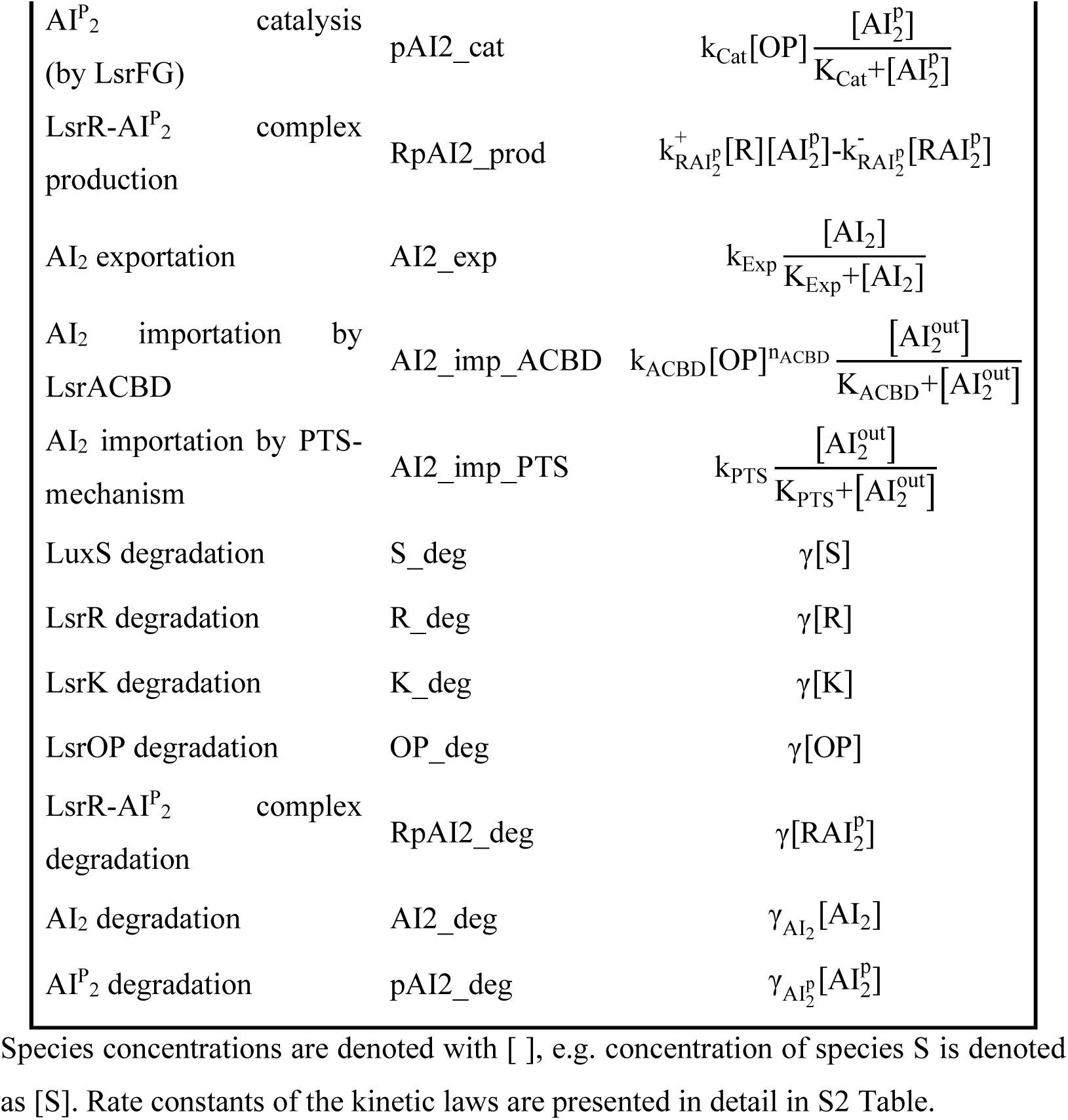
Kinetic laws of reaction network (QS module).

#### Kinetic model TTSS-1 module

We formulated a mathematical model (submodule) to account for the expression of TTSS-1. SPI1 genes are expressed hierarchically and exhibit a switch-like behavior transitioning from the “off” to the “on” state, as proposed in [35]. We constructed a minimal mathematical model (submodule) of the FA complex (InvF:SicA) positive feedback loop to quantify the dynamics of TTSS-1. The model describes a genetic circuit containing four proteins: the master regulator H (HilA), the transcription factor F (InvF), the chaperone A (SicA) and the secreted effector (translocon) C (SipC). The master regulator H regulates the expression of a large operon containing invF, sicA and sipC genes, and it acts as an intermediate between the microenvironment (i.e. environmental cues) and T3SS-1 module. A complex regulatory mechanism regulates H, as proposed in [35] and it will not be considered in this work.

Given that F and A regulators do not seem to affect H expression, as they mainly regulate the secreted effectors expression [48,49], we focus only on regulation of secretion assuming that the needle complex is already assemblied. To achieve this, we extended the network topology proposed by Temme *et al.* [36], descibed as the split (positive) feedback loop. In our approach though, we consider that the operon’s external promoter is up-regulated from H [50,51] and down-regulated from R, as discussed below. When expressed F bounds to chaperone A. FA complex upregulates another promoter, lying internally to the external promoter, from so on called internal promoter. The internal promoter controls the expression of the chaperon protein A and the effector protein C. Prior to the completion of functional needle, C binds to and sequesters A [52] forming AC (SicA:SipC) complex. Once the needle is functional, C is exported and A is free to bind F. FA then activates the internal promoter, thereby upregulating the expression of A and C proteins. The reaction network of the circuit is presented in Fig 2 (B) and the reactions’ kinetic laws in Table 2. Here, we assume that F and A and A and C form a 1:1 complexes. The synthesis of H, F, A, and C was each modeled by a single equation which contains protein synthesis, decay, cooperative binding, and repression. This mechanism leads to a system of 6 ODEs capturing the change in the total number of each protein in the cell. An interested user can find the system of ODEs and the corresponding rate parameters, in S1 and S3 Table, respectively. The details on the parameter estimation task is provided in Methods Section.

**Table 2.**
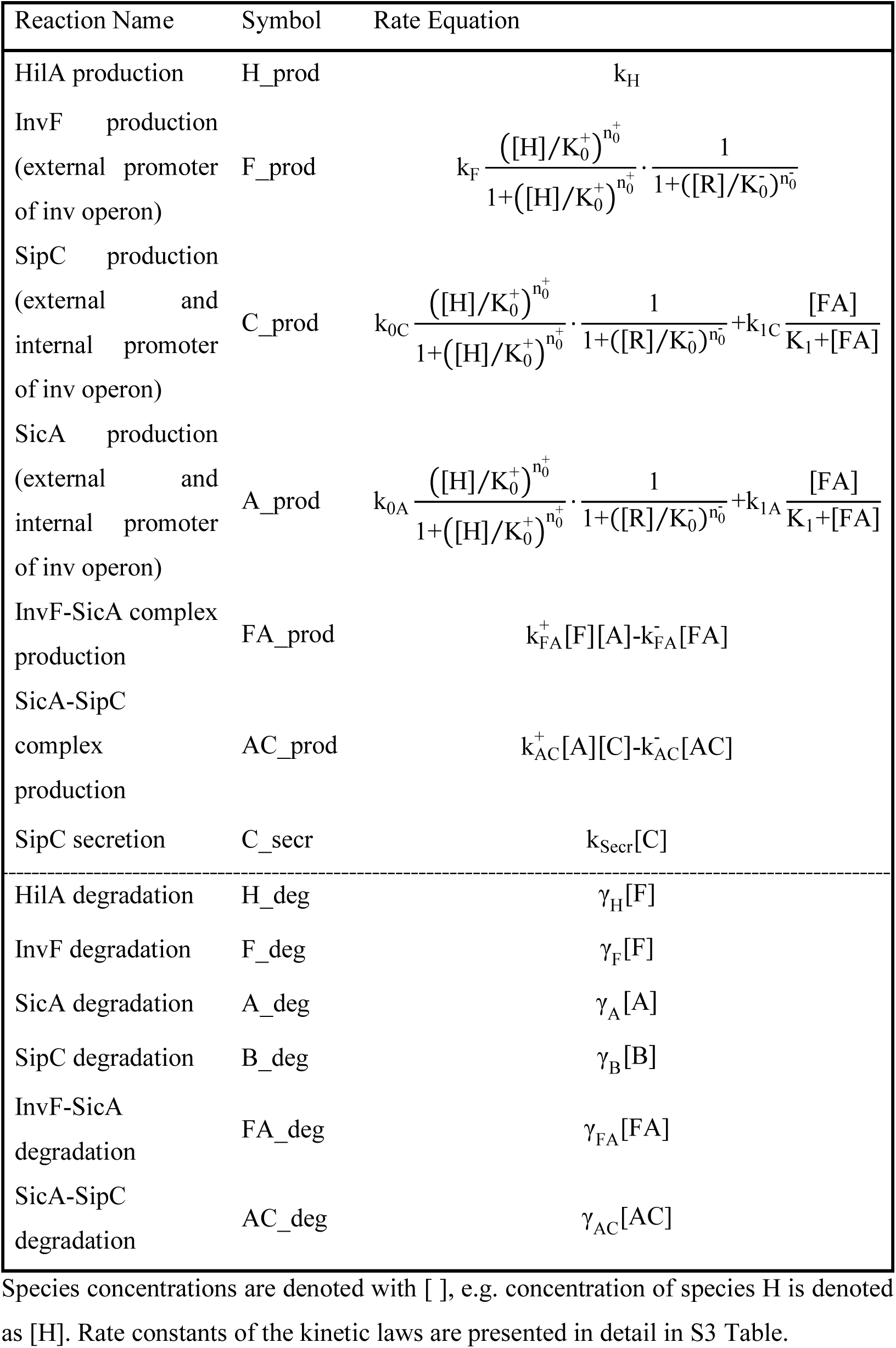
Kinetic laws of reaction network (TTSS-1 module).

| Reaction Name | Symbol | Rate Equation |
| --- | --- | --- |
| HilA production | H_prod | $k_H$ |
| InvF production<br>(external promoter<br>of inv operon) | F_prod | $k_F \frac{([H]/K_0^+)^{n_0^+}}{1+([H]/K_0^+)^{n_0^+}} \cdot \frac{1}{1+([R]/K_0^-)^{n_0^-}}$ |
| SipC production<br>(external and<br>internal promoter<br>of inv operon) | C_prod | $k_{0C} \frac{([H]/K_0^+)^{n_0^+}}{1+([H]/K_0^+)^{n_0^+}} \cdot \frac{1}{1+([R]/K_0^-)^{n_0^-}} + k_{1C} \frac{[FA]}{K_1+[FA]}$ |
| SicA production<br>(external and<br>internal promoter<br>of inv operon) | A_prod | $k_{0A} \frac{([H]/K_0^+)^{n_0^+}}{1+([H]/K_0^+)^{n_0^+}} \cdot \frac{1}{1+([R]/K_0^-)^{n_0^-}} + k_{1A} \frac{[FA]}{K_1+[FA]}$ |
| InvF-SicA complex<br>production | FA_prod | $k_{FA}^+[F][A] - k_{FA}^-[FA]$ |
| SicA-SipC<br>complex<br>production | AC_prod | $k_{AC}^+[A][C] - k_{AC}^-[AC]$ |
| SipC secretion | C_secr | $k_{Secr}[C]$ |
| HilA degradation | H_deg | $\gamma_H[F]$ |
| InvF degradation | F_deg | $\gamma_F[F]$ |
| SicA degradation | A_deg | $\gamma_A[A]$ |
| SipC degradation | B_deg | $\gamma_B[B]$ |
| InvF-SicA<br>degradation | FA_deg | $\gamma_{FA}[FA]$ |
| SicA-SipC<br>degradation | AC_deg | $\gamma_{AC}[AC]$ |

Species concentrations are denoted with [ ], e.g. concentration of species H is denoted as [H]. Rate constants of the kinetic laws are presented in detail in S3 Table.

In our kinetic model we combine LuxS-QS and TTSS-1 modules via R repressor. In [38], Choi *et al.* report that R, negatively controls the expression of several TTSS-1 proteins. Specifically R controls negatively F, A and C expression as proposed in [38], so by considering this finding we assume multiple trancription factor regulatory mechanism on the external promoter of the large operon, see the kinetic laws of F, A, and C production reactions in Table 2 for details. In order to examine, each reaction individually, we provide the set of reactions, see S1 Fig, corresponding to the proposed reaction network.

#### Ibm

The proposed reaction network of LuxS-QS and TTSS-1 interconnected activity was incorporated into an individual-based modeling simulation. Specifically, we developed a mixed cell-based/ODE-model which captures the molecular machinery of cells and the mechanical interactions between them. In this model, we describe bacteria as two half-spheres connected by a cylinder. Cells grow in the direction of the cylinder and divide perpendicular to this direction. We model mechanical interactions explicitly so as to minimize spatial overlap in the colony, see in the Methods section for more details. In this approach, intracellular AI_2_ activity depends on its production and degradation by the cell and on the cell wall permeability as well as on the diffusive properties of the microenvironment’s medium. In order to capture this dependency, we extended the kinetic model’s reactions network (see in Fig 2 (A) the species and interactions colored in gray). Specifically, we added two reactions; one for S (LuxS) synthesis [53] and another for AI_2_ production by S inside cell. Intracellularly produced AI_2_ is then externalized to then environment by an unknown mechanism, as proposed in [11]. In Table 1, we provide the kinetic laws for each added reaction. The parameters of these reactions can be found in S2 Table. An interesting reader can find the multicellular system of ODEs in S1 Table. Furthermore, we assume that 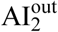 diffuses freely in the environment. In our model bacteria growing exponentially, we take also into account the effect of dilution on species concentrations. In the Methods section, we describe thoroughly the bacterial growth and molecule diffusion of the proposed modelling approach.

### Model dynamic behavior

#### Kinetic model

To illustrate the dynamic behavior of the proposed kinetic model, we used the COPASI simulation software [54] using specifically the LSODA algorithm [55]. Below we show the results of model simulation, which last 30 “lab” hours.

In Fig 3, we present the dynamic behavior of the proposed (kinetic) model. The model is stimulated by two different pulses. The dark blue pulse simulates AI_2_ while the green pulse simulates the activation and deactivation of H in the micro-environment.

**Fig 3.**
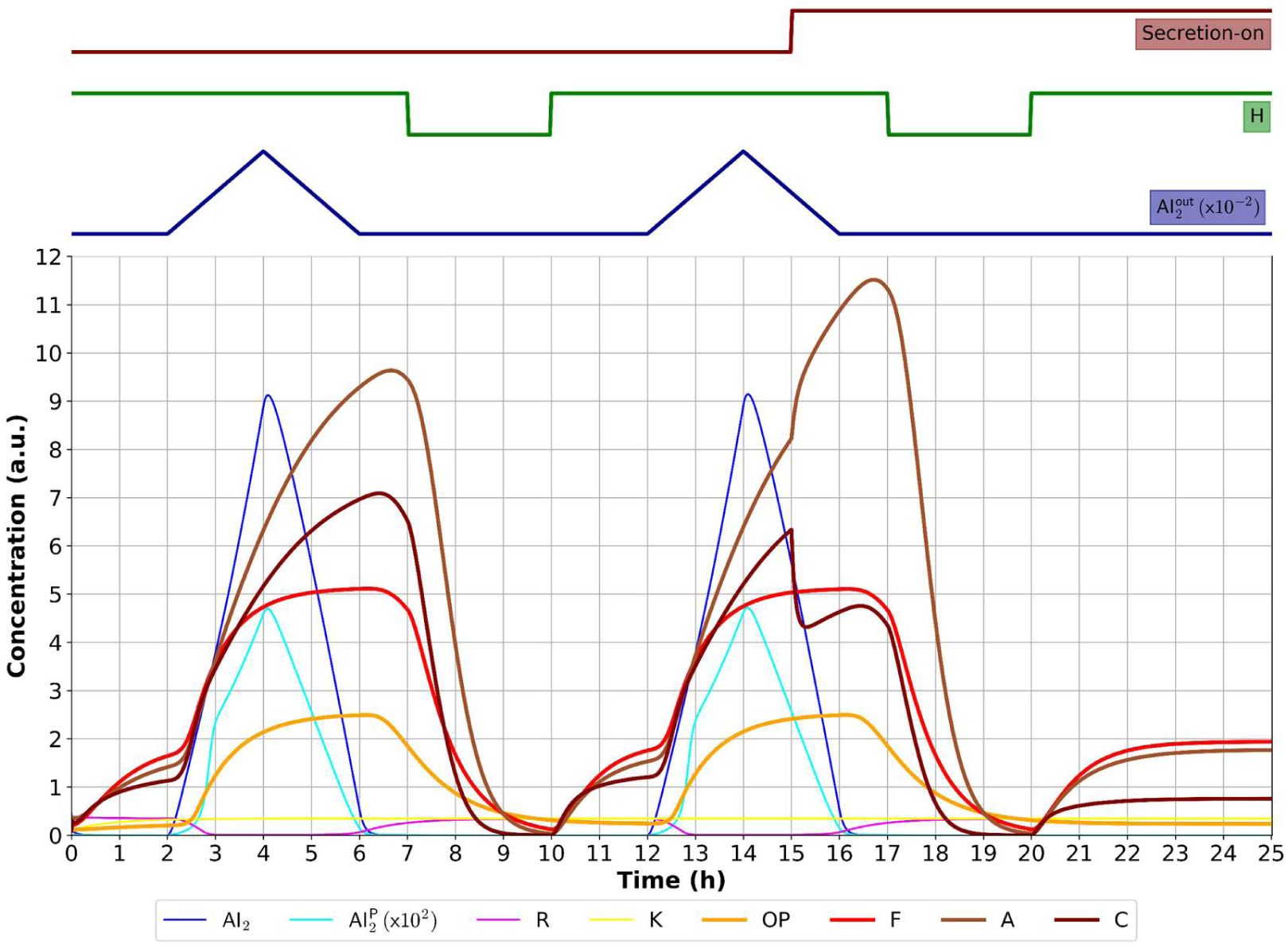
Dynamic behavior of the proposed kinetic model. Colors represent the different species of the model. Dark blue input pulse simulates 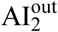 concentration in the micro-environment. Green input pulse simulates H activation and deactivation, i.e. virulence inducing and non-inducing conditions in the micro-environment. The rest of the species represent model’s outputs. See text for details.

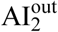 is introduced to the microenvironment, between 2 and 6 hours (blue pulse). Simultaneously, H is triggered between 0 and 7 hours (simulating virulence inducing conditions). At 2 hours, 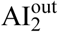 is imported to the cell (denoted as AI_2_) and K starts phosphorylating it. R stops repression of lsrRK, lsr operon and TTSS-1 species because it binds to 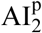. TTSS-1 species are upregulated due to the existence of H. Within four hours, AI_2_ reaches maximum expression. F starts associating with A and the FA complex boosts the production of A and C reaching their maximal expression between 6 and 7 hours. The presence of the positive feedback loop yields the observed delay between AI_2_ and R repressed species maximal expression. After 6 hours AI_2_ is depleted from the cell causing all R repressed species to decrease dramatically. At 7 hours, H is deactivated, resulting to zero TTSS-1 species. We must notice that if H is activated when AI_2_ is depleted from the cell (at 24 hours), TTSS-1 species are expressed in reduced level. Furthermore, we observe that even if the pulses’ pattern is the same between 12 and 19 hours, the TTSS-1 species follow similar but not the same trends as in previous interval. This happens because we induced secretion of C to the simulation, i.e. we enabled secretion of SipC at 17 hours (t_secr_). At this time, H was activated. At 19 hours, H was deactivated, in order to observe the relaxation dynamics of the model. The time point t_secr_ was selected in a way to allow C to reach its steady state both before and after the triggering of secretion. In summary, the model can be led to 4 distinct states (with or without secretion), according to the input pulses. In Table 3, we present an overview of the different states of the proposed model.

**Table 3.**
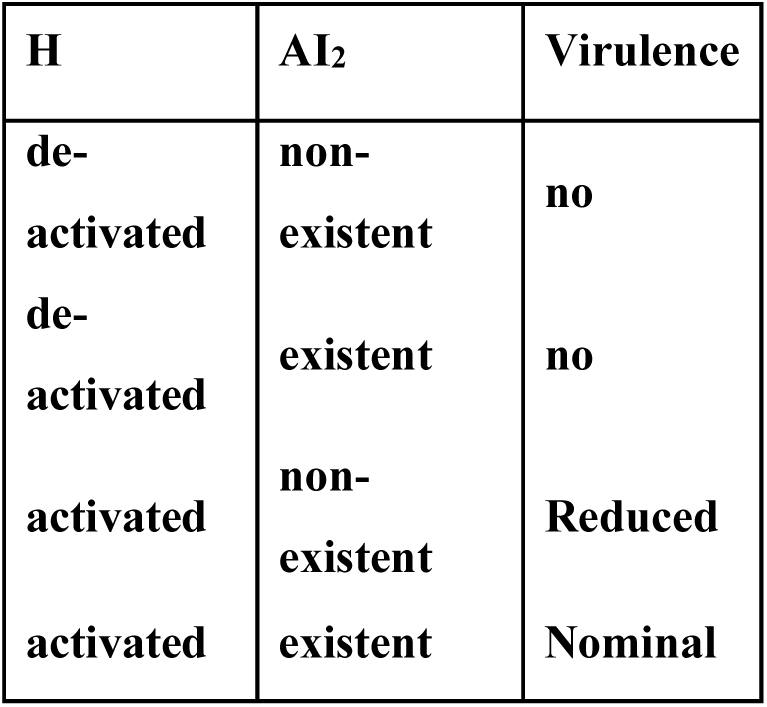
Different states of the proposed model.

| H | $AI_2$ | Virulence |
| --- | --- | --- |
| de-activated | non-existent | no |
| de-activated | existent | no |
| activated | non-existent | Reduced |
| activated | existent | Nominal |

The behavior of OP is consistent with the experimental findings of Taga *et al.* [46]. The LuxS-QS feedback loop is reproduced successfully in-silico even with a parsimonious model as far as QS species are concerned. Furthermore, the behavior of A is consistent with experimental findings of Temme *et. al* [36] even if we recapitulate what they proposed with an extended model that combines LuxS-QS feedback loop with the TTSS-1 “split” positive feedback loop. Specifically, A continues to be expressed for 100 min even in non-inducing conditions (H was deactivated 7 hours) and it is also highly expressed when secretion is activated as proposed in [36]. However, in our model for both cases, AI-2 expression is a prerequisite for virulence activation because it relieves TTSS-1 species repression from R.

#### Ibm

To illustrate the dynamic behavior of the proposed Ibm, we used the Cellmodeler simulation software [56] using specifically Crank-Nicholson algorithm [57]. Below we show the results of model simulation, which last 12 “lab” hours.

Here, we present the dynamic behavior of Ibm. compared with the behavior of kinetic model. As discussed in model development subsection, in this modelling approach each cell expresses S constantly (until 4 hours), and thus, it produces AI_2_ which is in turn exported to the micro-environment. Even though we do not trigger the model externally (with 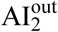), we obtain similar dynamic behavior on average as in kinetic model, see Fig 4 (A1) and (A2). To compute the average concentration of each species per time point, we follow the formula below:

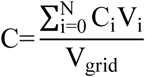

**Fig 4.**
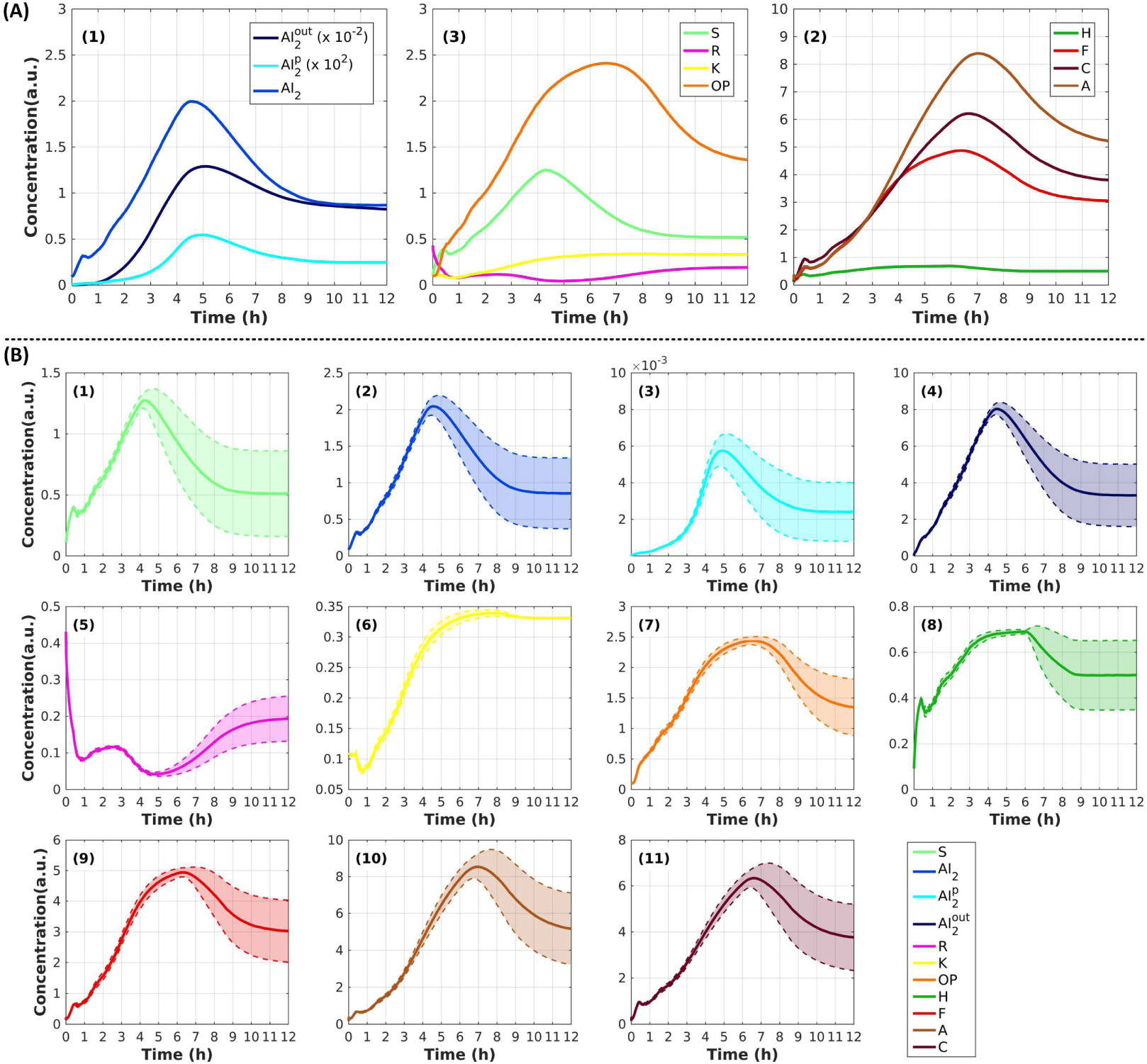
Dynamic behavior of Ibm. Colors represent the different species. (A) On average species expression: (1) Autoinducer species, (2) LuxS-QS species, and (3) TTSS-1 species. We notice that species activity follows the same trend as in kinetic model simulation between 0 to 12 hours, thereby validating our Ibm approach. (B) Noise in growth leads to different AI_2_ pulses for each cell and thus to variability in species expression. LuxS-QS species variability increases rapidly after 4 hours (experimental time) while TTSS-1 species increases rapidly after 6 hours.

where C is the species concentration, V_grid_ is the grid volume, C_i_ is the species concentration of a cell and V_i_ is the volume of a cell.

In Fig 4 (B), we can observe the heterogeneity of each species of the simulation. Our model recapitulates the stochasticity existing in all the biological phenomena; this is the potential of individual-based modeling. Ibm approach enabled us to drill down to single-cell level and examine how exposure of each cell to AI_2_ can lead in different species expression in a clonal population. It is obvious that stochasticity plays a significant role at population level. Each cell contributes in a different fashion to the whole population virulence phenotype.

### Model validation

Following the model development and its dynamic behavior analysis, we validate it in terms of reproducing the behavior (qualitatively) of the LuxS-QS and TTSS-1 species on the basis of experimental data found in the literature. It should be mentioned that those experimental data were not considered in the parameter estimation procedure, i.e. in model calibration. Furthermore, it is of paramount importance that all the knock-out experiments presented herein produce similar results in both modelling approaches. Here, we present the simulated results produced either by the kinetic model or by the Ibm depending on the case study. In case the results of the case study can be produced by both the kinetic model and the Ibm, then we provide here the results produced by the kinetic model and we provide the corresponding results of the Ibm in S1 Text.

#### AI_2_ dynamics [46]

First, we evaluate if our model reproduces the autoinducer dynamics inside and outside cell. This case study can only be examined by the Ibm which is able to capture the dynamics of every autoinducer species variation in the micro-environment, i.e. AI_2_, 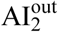 and 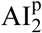,). In Fig 4 (A1), we illustrate how autoinducer species evolve in time during the simulations. It can be observed that model’s species follow the same trend as the species of the kinetic model approach and have approximately equal level of expression on average. Our results, on external 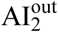 and 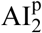 species evolution are confirmed by *Taga et al.* [46], see Fig 3 (B) and (D) in their manuscript.

Then, we did knock-outs experiments of *luxS* and lsr genes in silico and we monitor how they affect lsr operon expression, see Table 4 and Table 5 for kinetic model results and S2 Fig for the corresponding Ibm results.

**Table 4.**
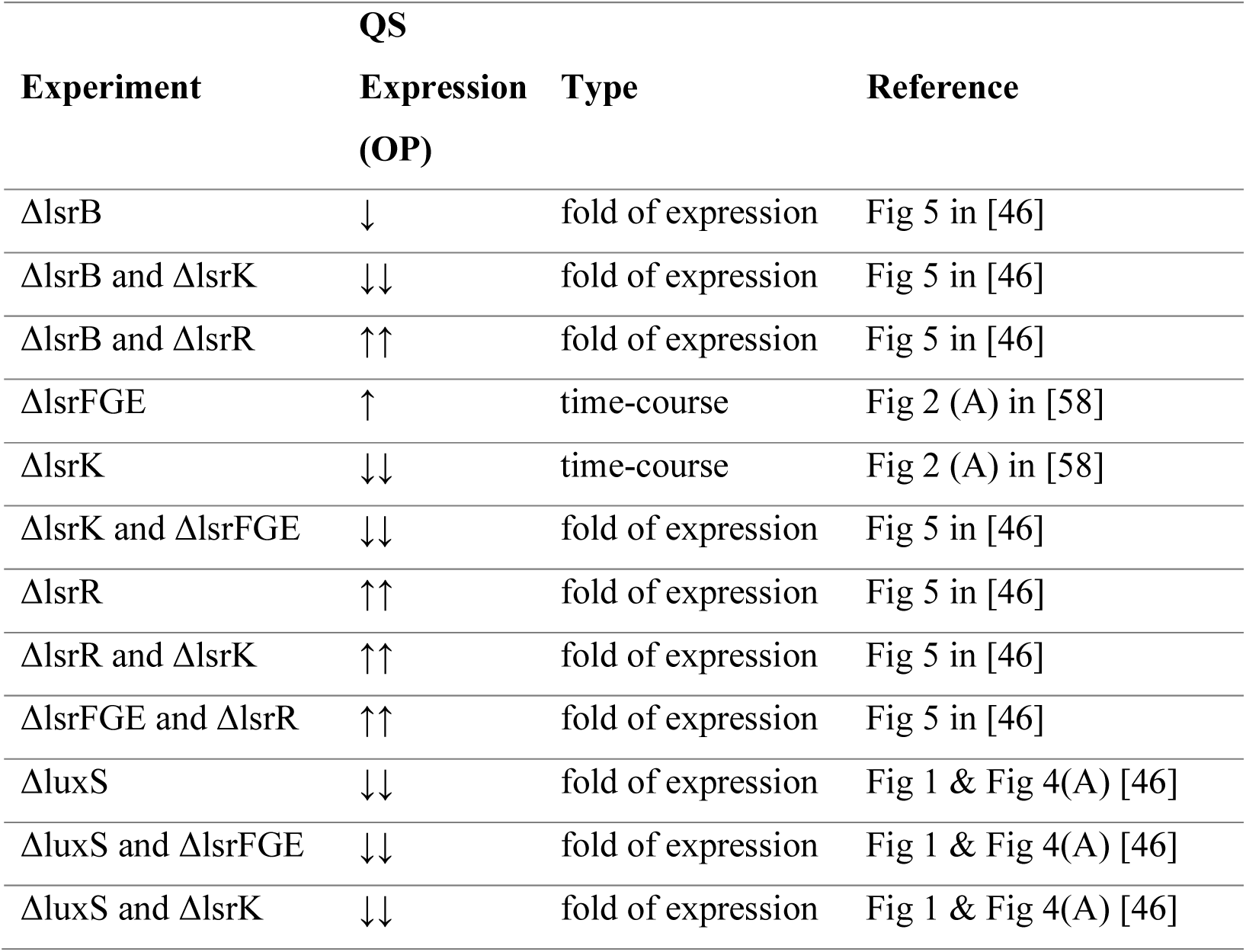
In silico reproduction of wet lab experiments affecting QS expression.

| Experiment | QS |  | Reference |
| --- | --- | --- | --- |
|  | Expression (OP) | Type |  |
| ΔlsrB | ↓ | fold of expression | Fig 5 in [46] |
| ΔlsrB and ΔlsrK | ↓↓ | fold of expression | Fig 5 in [46] |
| ΔlsrB and ΔlsrR | ↑↑ | fold of expression | Fig 5 in [46] |
| ΔlsrFGE | ↑ | time-course | Fig 2 (A) in [58] |
| ΔlsrK | ↓↓ | time-course | Fig 2 (A) in [58] |
| ΔlsrK and ΔlsrFGE | ↓↓ | fold of expression | Fig 5 in [46] |
| ΔlsrR | ↑↑ | fold of expression | Fig 5 in [46] |
| ΔlsrR and ΔlsrK | ↑↑ | fold of expression | Fig 5 in [46] |
| ΔlsrFGE and ΔlsrR | ↑↑ | fold of expression | Fig 5 in [46] |
| ΔluxS | ↓↓ | fold of expression | Fig 1 & Fig 4(A) [46] |
| ΔluxS and ΔlsrFGE | ↓↓ | fold of expression | Fig 1 & Fig 4(A) [46] |
| ΔluxS and ΔlsrK | ↓↓ | fold of expression | Fig 1 & Fig 4(A) [46] |

**Table 5.** In silico reproduction of wet lab experiments affecting virulence expression.

| Experiment | Virulence |  | Reference |
| --- | --- | --- | --- |
|  | Expression Species | Type |  |
| ΔinvF | ↓ | fold of expression | Fig 1(A) & Fig 2(A) [38] |
| ΔluxS | ↓↓ | fold of expression | Fig 1(A) & Fig 2(A) [38] |
| ΔinvF and ΔluxS | ↓↓ | fold of expression | Fig 1(A) & Fig 2(A) [38] |
| ΔlsrR | ≈ | fold of expression | Fig 2(A) [38] |
| ΔluxS and ΔlsrR | ≈ | fold of expression | Fig 2(A) [38] |
| *lsrR*<sup>+</sup> plasmid | ↓↓ | fold of expression | Fig 3 in [38] |
| $\Delta\text{sicA}$ | $\downarrow\downarrow$ | fold of expression | [36] |

#### Mutations to the SicA:InvF-binding site (internal promoter) [36]

To simulate this mutations in silico we varied the binding constant (K) that has a significant effect on the strength of the positive feedback loop of TTSS-1 module, as claimed in [36]. Specifically, we varied K over a veritable range of values, from 3.92 to 7.84 (with step 0.3922) under non-inducing conditions, i.e. H expression de-activation. In Fig 5 (A), we present the results of our simulations confirming that the persistence of A expression can be varied giving a remarkable diversity of relaxation times, as shown in [36] Fig 9 (b). Furthermore, we trigger secretion in our model (at 5 hours) and we observe that the expression of A is amplified, i.e. the activity of the feedback loop increases and the persistence of A expression is higher in comparison to the feedback loop without secretion, see Fig 5 (B). This finding, it is also shown in in [36] Fig 8 top left. It is noteworthy that our model recapitulates the findings of [36] having also integrated LuxS-QS mechanism regulating the TTSS-1 feedback loop. The same results are obtained by our Ibm approach, see S4 Fig.

**Fig 5.**
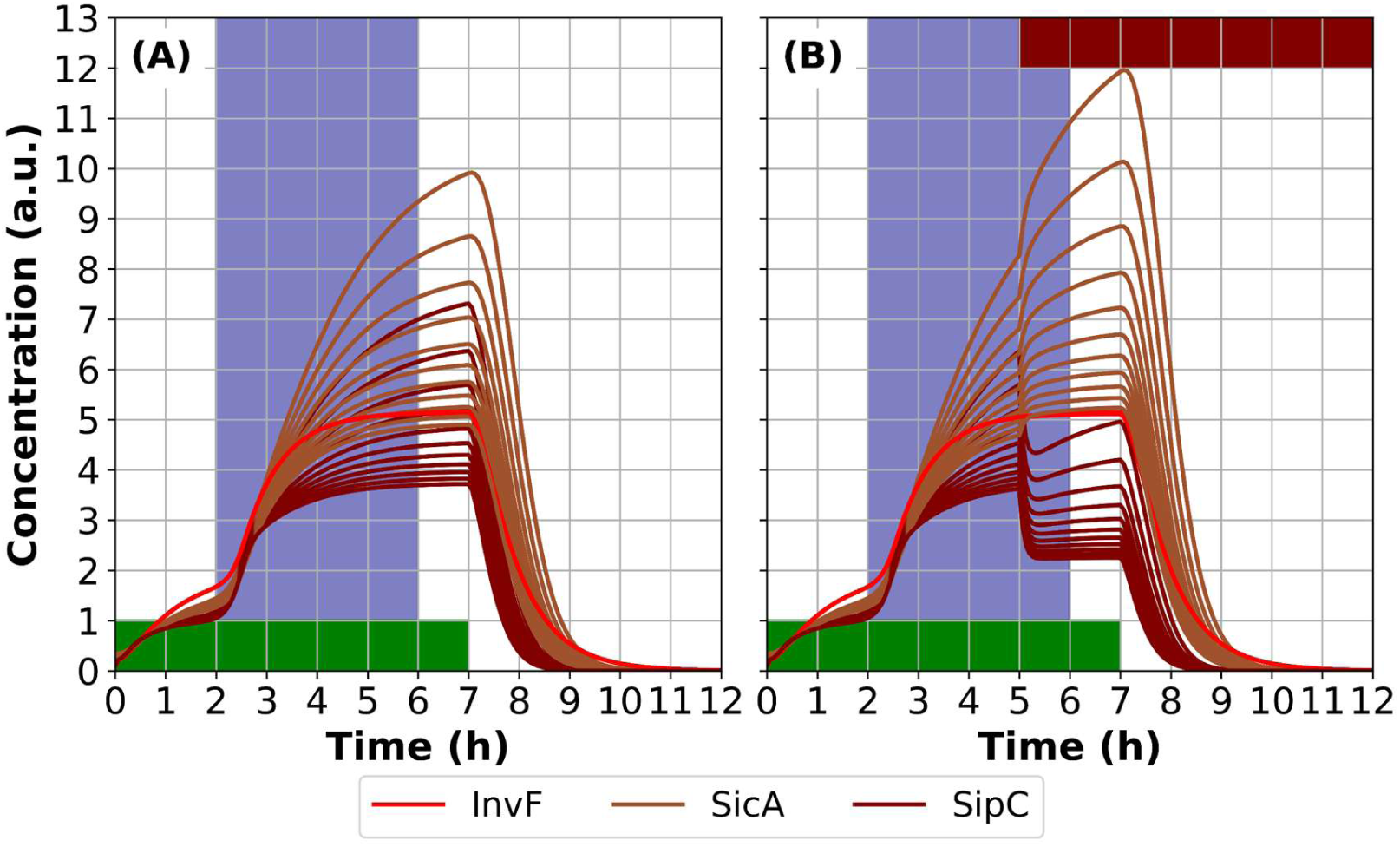
Mutations to the SicA:InvF binding site effect on A expression. Colors represent the different species. Dark blue region represents the input 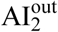 pulse. Green region represents the interval that H is activated. Marron region represents the interval that secretion is activated. Time-course results for 12 hours experimental time. Notice that when binding constant (K) increases, A expression decreases leading to lower relaxation times both in secreting and non-secreting mechanism. Here we varied K from 3.92 to 7.84 (with step 0.3922).

#### Ιbm recapitulates the bistability of TTSS-1 expression [36,58]

Under inducing conditions, single-cell reporters for TTSS-1 species indicate that cells can be found in “on” or in “off” state. TTSS-1 expression is induced by growth-related environmental signals (i.e. autoinducer molecules) as reported in [58]. In our approach, the regulatory mechanism controlling TTSS-1 expression in each cell is induced due to micro-environment stressors (by H activation/deactivation as previously discussed) and LuxS-QS community effect. Our model introduces noise in growth leading to different LuxS-QS expression patterns which in turn lead to different TTSS-1 expression in S. *Typhimurium* clonal population under inducing conditions, see Fig 6 (A)-(C). We can observe that the community’s spatial distribution plays a significant role in the expression of TTSS-1, in the sense that, single-cell virulent phenotype depends on the exposure of each cell to 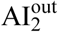 availability. In Fig 7 (D), it can be observed that the initiation rate of TTSS-1 expression seemed to increase upon entry into the late logarithmic growth phase approximately after 3.5 hours, as shown in [58] Fig 3 (D). Such behaviour can be reproduced in-silico only by Ibm approaches.

**Fig 6.**
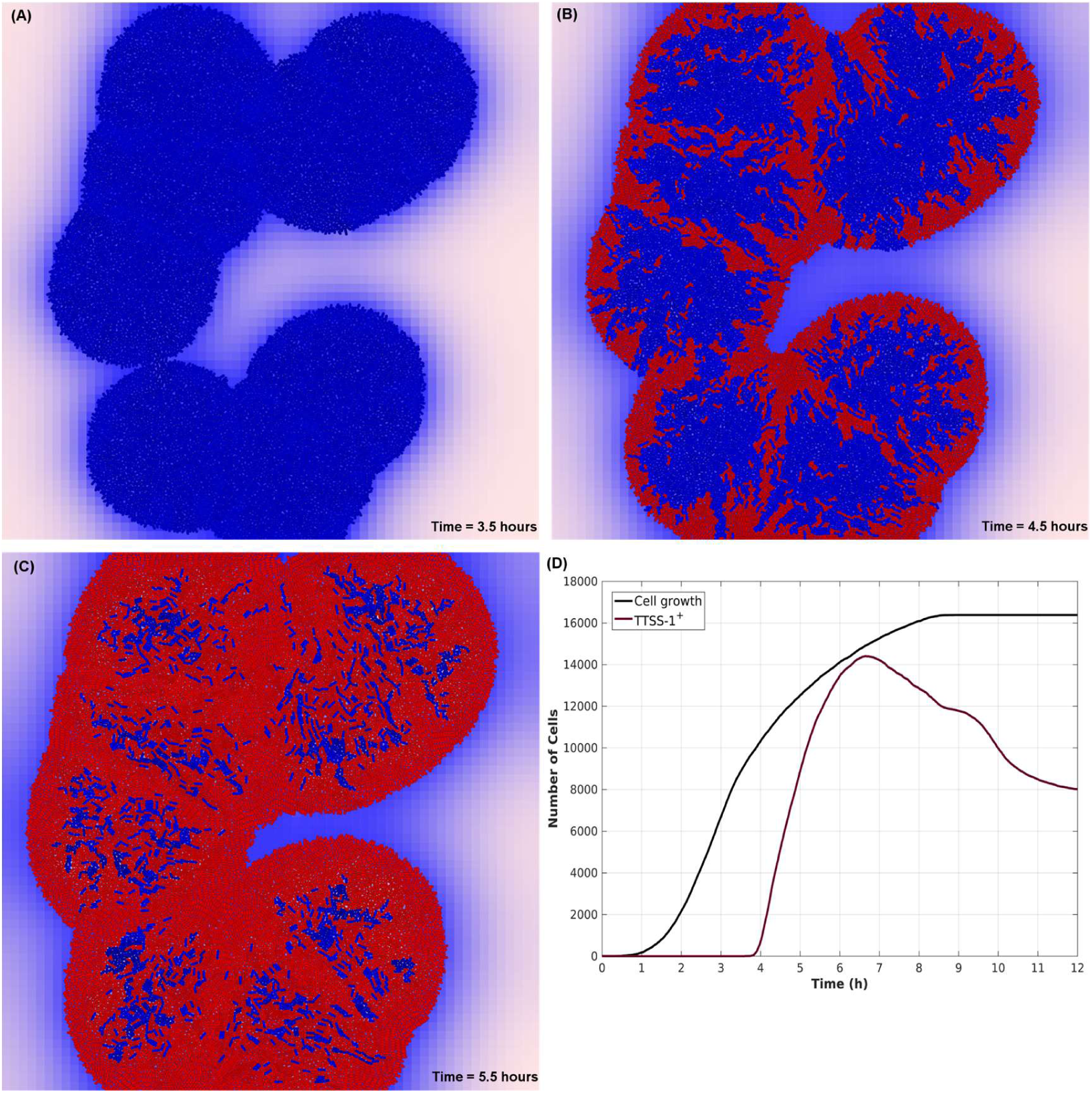
Topological distribution of TTSS-1 cells. Simulation for 9 hours experimental time. Virulence inducing conditions between 0-6 hours. **(A)-(C)** Blue color represent AI_2_ expression in each cell. Maroon color represents A expression in each cell. Blue color represents 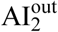 activity in the micro-environment. The higher the color intensity the higher the species expression. Cells become maroon when TTSS-1 mechanism is activated. Different patterns of TTSS-1 expression depend on the exposure of each cell to 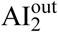 concentration and to stochastic growth. **(D)** Growth curves of the population and TTSS-1^+^ subpopulation with and without secretion. We can observe that virulent subpopulation start rising when entering in the late exponential phase of growth, as claimed in [58]. Furthermore, our model predicts that a TTSS-1^+^ secreting (virulence factors) subpopulation is more aggressive that a non-secreting subpopulation.

**Fig 7.**
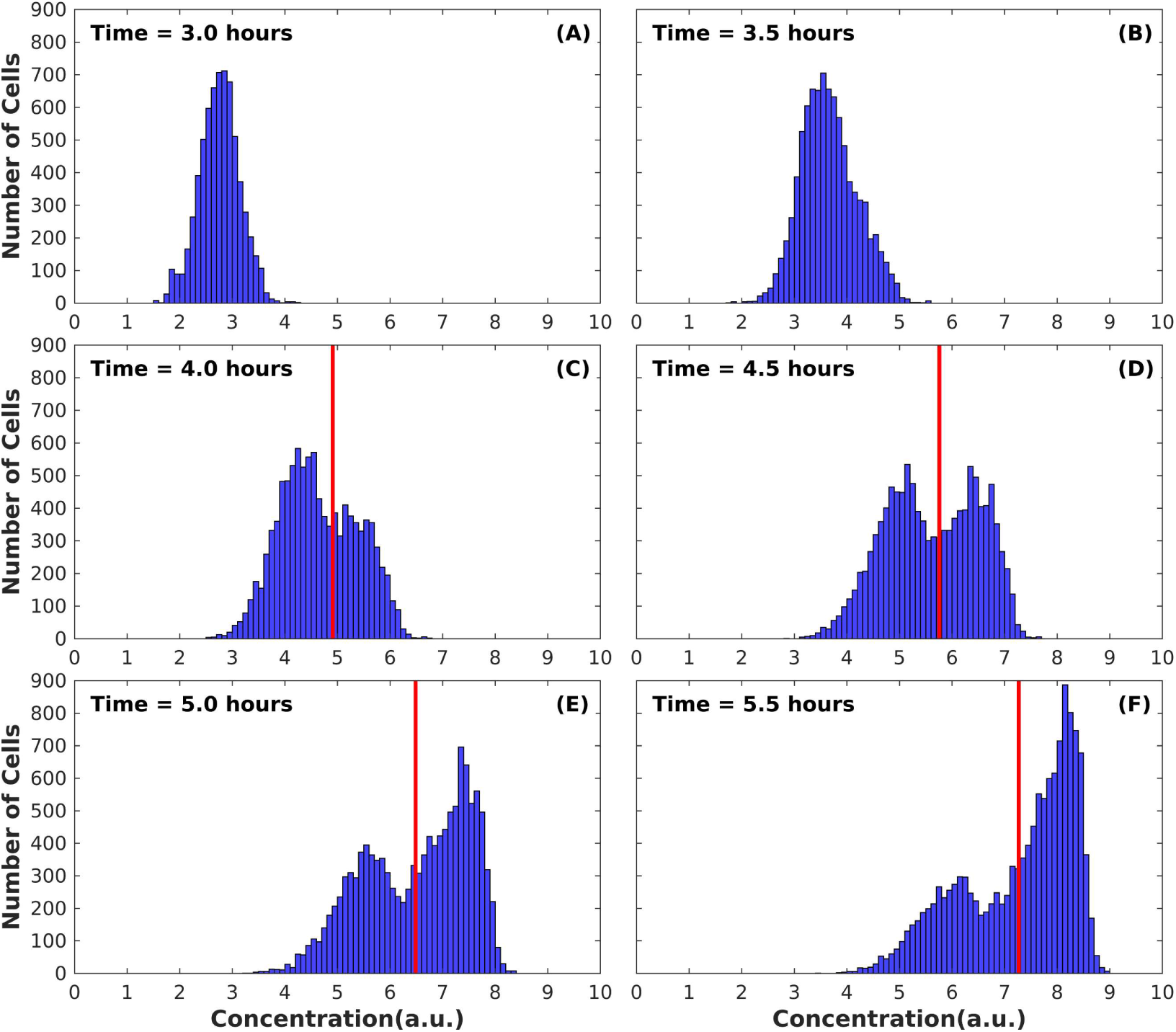
Ibm recapitulates bistable expression of A in S. *Typhimurium* microbial communities. Simulation for 12 hours experimental time. Virulence inducing conditions between 0-6 hours. A distribution evolution at population level. After 4 hours of growth (late exponential phase see Fig 6 (D)), the distribution becomes bimodal. TTSS-1^+^ subpopulation emerges from the whole population. Red lines are provided to enable visual inspection of the two distributions modes. Notice that the distribution average increases in time, but the bimodal pattern insists.

To further support that the dynamic interplay of TTSS-1^+^ and TTSS-1^−^ individuals in a clonal *S. Typhimurium* population depends on the stochasticity introduced in cell growth, via LuxS-QS, we provide the bistable A expression evolution in time, see Fig 7. Our findings are similar with the distributions that are experimentally produced in [36] Fig 3 SicA expression under virulence induction. In S1 Movie, we provide a simulation that fully captures bistability dynamics of the proposed approach. It is noteworthy, that our model recapitulated bistability in TTSS-1 expression without further calibration than the initial.

#### Slow growing cell are more appropriate to become TTSS-1^+^

Our Ibm suggests that stochasticity in growth leads to a fast and a slow growing subpopulation. And thus, in virulence inducing conditions the slower subpopulation becomes the TTSS-1^+^ subpopulation while the faster becomes the TTSS-1^-^. In [58], Sturm *et al.* analyzed the fitness costs associated with the expression of TTSS-1. They proved experimentally that TTSS-1^+^ cells had a reduced growth rate. So, they claimed that TTSS-1 expression represents a burden for the growth of single-cells. Inspired by this finding, we managed to recapitulate this behavior in silico (without further tuning the initial model). Thus we conlcuded that in a virulence inducing micro-enviroment, stochasticity in growth may lead to slower subpopulation that it is more “appropriate” to become TTSS-1^+^, see Fig 8. Intuitively, this finding makes sense, taking into account the bet-hedging strategy follow by bacteria, however it remains to be proved experimentally.

**Fig 8.**
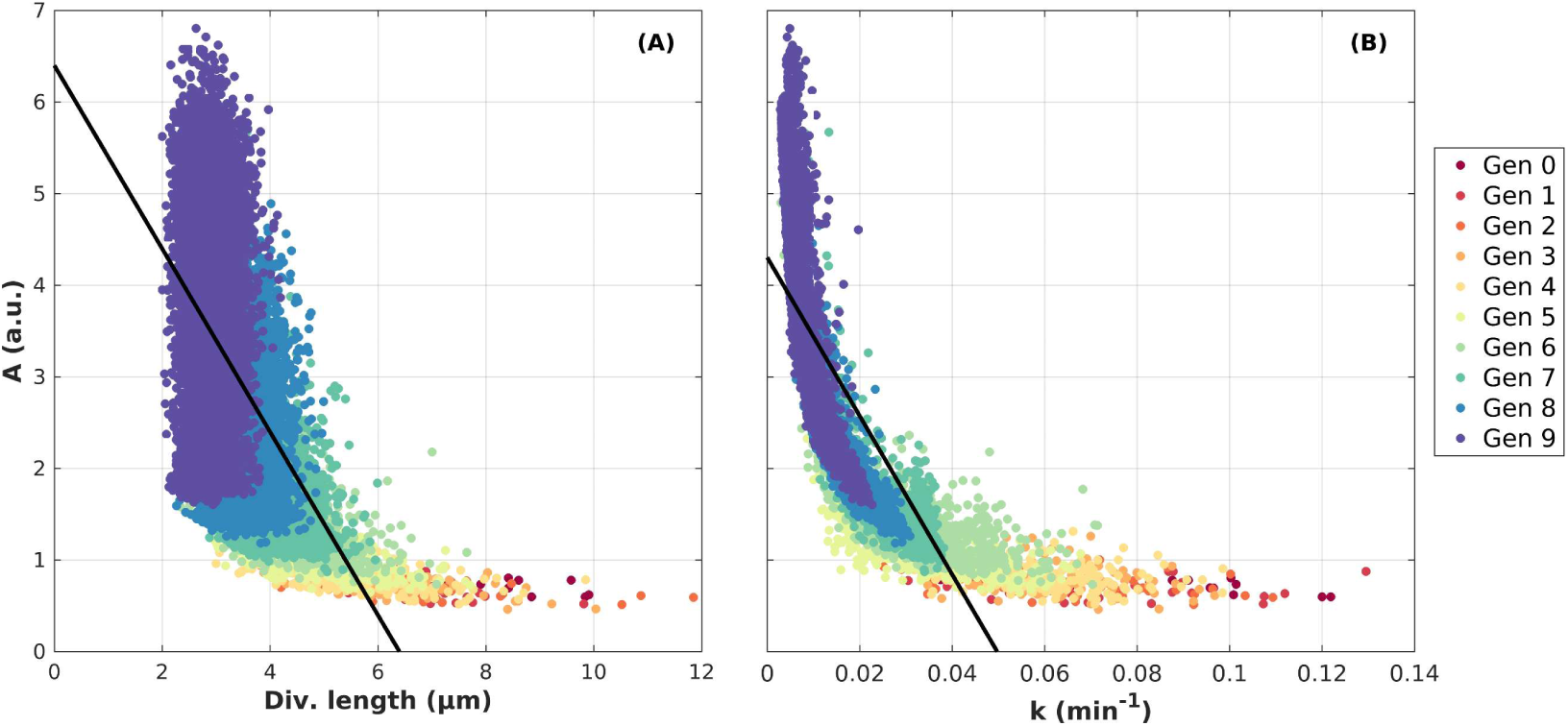
Growth impact on TTSS-1^+^ phenotype. (A) Scatterplot of A expression vs Division length. (B) Scatterplot of A expression vs *k* growth rate. Different colors represent different generations of the cell population. Both division length and *k* growth rate are negatively correlated with expression of A.

### Model predictions

The developed model was used as a testing platform for many hypothetical scenarios. Although no quantitative information are available for our model species to compare to, it is still possible to draw some conclusions regarding LuxS-QS and TTSS-1 mechanisms. It is also important to note that all hypotheses were tested by changing the relevant parameters of the system and not any of the initial conditions. Based on the “nature” of the system, the long-term goal would be to unveil possible experimental intervention strategies aiming at TTSS-1 via tuning LuxS-QS.

As a first testing scenario, our model is set to predict the expression of TTSS-1 species in LuxS-QS gene knock-out experiments. The results produced by the kinetic model are presented in Table 6, while the corresponding results of the Ibm are provided in S5 Fig.

**Table 6.** In silico wet lab experiments predicting virulence expression.

| Experiment | Virulence |  |
| --- | --- | --- |
|  | Expression | Type |
|  | Species |  |
| ΔlsrB | ↓ | fold of expression |
| ΔlsrB and ΔlsrK | ↓↓ | fold of expression |
| ΔlsrB and ΔlsrR | ≈ | fold of expression |
| ΔlsrFGE | ≈ | fold of expression |
| ΔlsrK | ↓↓ | fold of expression |
| ΔlsrK and ΔlsrFGE | ↓↓ | fold of expression |
| ΔlsrK and ΔlsrR | ≈ | fold of expression |
| ΔlsrR and ΔlsrFGE | ≈ | fold of expression |

#### 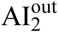 amplitude variation

In this work, we argue that LuxS-QS could act as a “tuning knob” which controls how long A expression persists after the TTSP-1 is deactivated. To test this assumption, we varied this 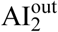 amplitude into a physiologically relevant range in non-inducing virulence conditions. Our model predicts that the virulence persistence duration can be varied by varying the strength of the positive feedback loop, resulting in a remarkable diversity of relaxation times with or without secretion (see Fig 9). Our model provides similar relaxation patterns with those generated by varying binding constant (K), as discussed in model validation subsection (Fig 7). It is remarkable that LuxS-QS mechanism when tuned accordingly is apt to produce a result that has previously been claimed to be achieved only by genetic mutations, as claimed in [36]. The TTSS-1 positive feedback loop can be tuned by LuxS-QS variation except for mutations to the SicA:InvF-binding site (internal promoter).

**Fig 9.**
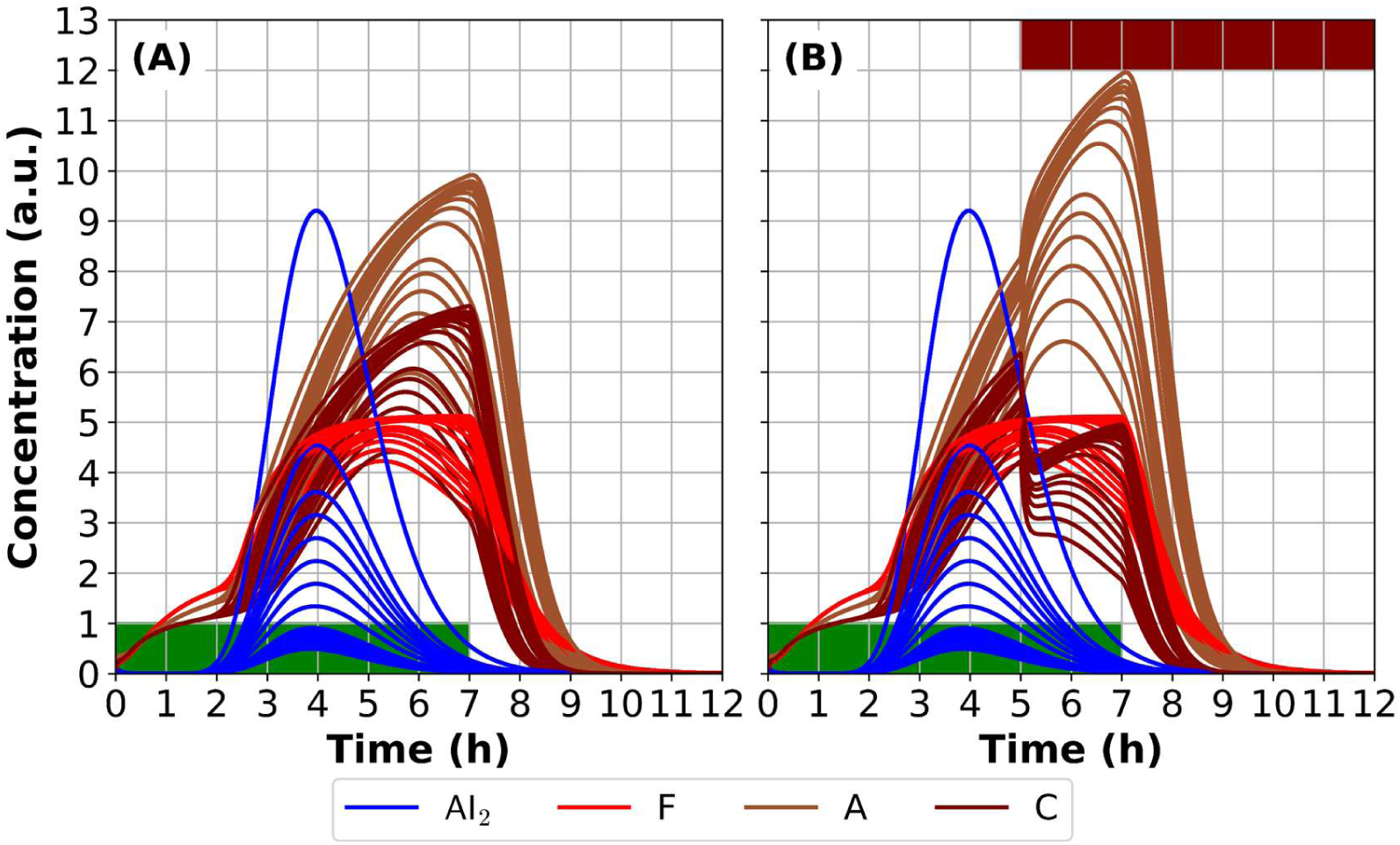
Amplitude of 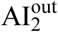 pulse variation leads to different relaxation times of A. Colors represent the different species. Green region represents the interval that H is activated. Time-course results for 12 hours experimental time. When AI-2 amplitude increases, A expression increases too leading to higher relaxation times. Here we varied AI-2 amplitude from 0.006 to 0.05. Notice that we do not alter the value of Binding constant (K), as we did in Model validation subsection.

Furthermore, in individual-based modelling approach, we attained indirectly the same results. Stochasticity in growth effects AI_2_ production in each cell, while simultaneously cell’s exposure to 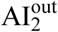 is different depending on its position, so each cell has different relaxation time of A. In Fig 4 (B), we can observe, that variance in AI_2_ species leads to variance in virulence persistence through A.

### A digital twin of Nanofactories triggering anti-cancerous response in targeted bacteria

The proposed approach was inspired by the experimental work presented in [8,9]. Our purpose is to show that TTSS-1 virulence module, which is a killing weapon for normal cells [59], can be expressed on demand and synthesize and secrete effectors, converting it to a killing weapon for cancerous cells [20,60]. This represents a new type of controller for targeted drug delivery as actuation (synthesis and delivery) depends on a receptor density marking of the diseased cell.

Exploiting, the knowledge acquired from the calibration of the kinetic model and its application on Ibm, we decided to proceed to a realistic experiment of synthetic biology nature. Initially, we modified our model to simulate programmed motility based on the concentration of 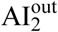 in the micro-environment. In that respect, we were able to direct bacteria motility to 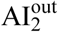 gradient produced by nanofactories (NF) [64]. In silico, we design NFs that can dock on human epidermal growth factor receptor 2 (HER2) [61] (HER2 are found on colorectal cancer cells (CRC) [62]). AI_2_ production depends on the on-site density of HER2. So, when cells reach CRCs, they initiate the expression of TTSS-1 based on HER2 density.

In Fig 10, we describe the simulation setup. The simulation comprises of anti HER2 nanofactories (anti-HER2-NF) (blue circular objects in the right side of each panel in Fig 10) docked onto mammalian cell surfaces, synthesizing AI_2_ signal. In the specific setup we assume that HER2 are uniformly distributed on the cell surface. The docking of anti-HER2-NF onto the surfaces is analogous to HER2 density which, in turn, controls the subsequent AI_2_ synthesis, bacteria migration, and the switching response phenotype. The motile bacteria (initially lying in the left side of Fig 10 (A)) have the ability to express TTSS-1 species but not S. Furthermore, motile bacteria sense 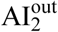 and swim towards maximum gradient of 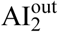. When they reach the source of AI_2_ production, i.e. NFs onto cancerous cells, they start growing and expressing their virulent phenotype (Fig 10 (B)-(I)), given that there is high-osmolarity and low oxygen in the micro-environment, i.e. simulating micro-environment of intestinal epithelium [63,64]. The efficiency of the system is quantified via the expression of the translocon protein C (which could play the role of translocator of a surrogate drug to the cancerous cells). For example, in the specific setup, C is highly expressed (> 3 a.u.) in some cells after 7 hours (see Fig 10 (D)).

**Fig 10.**
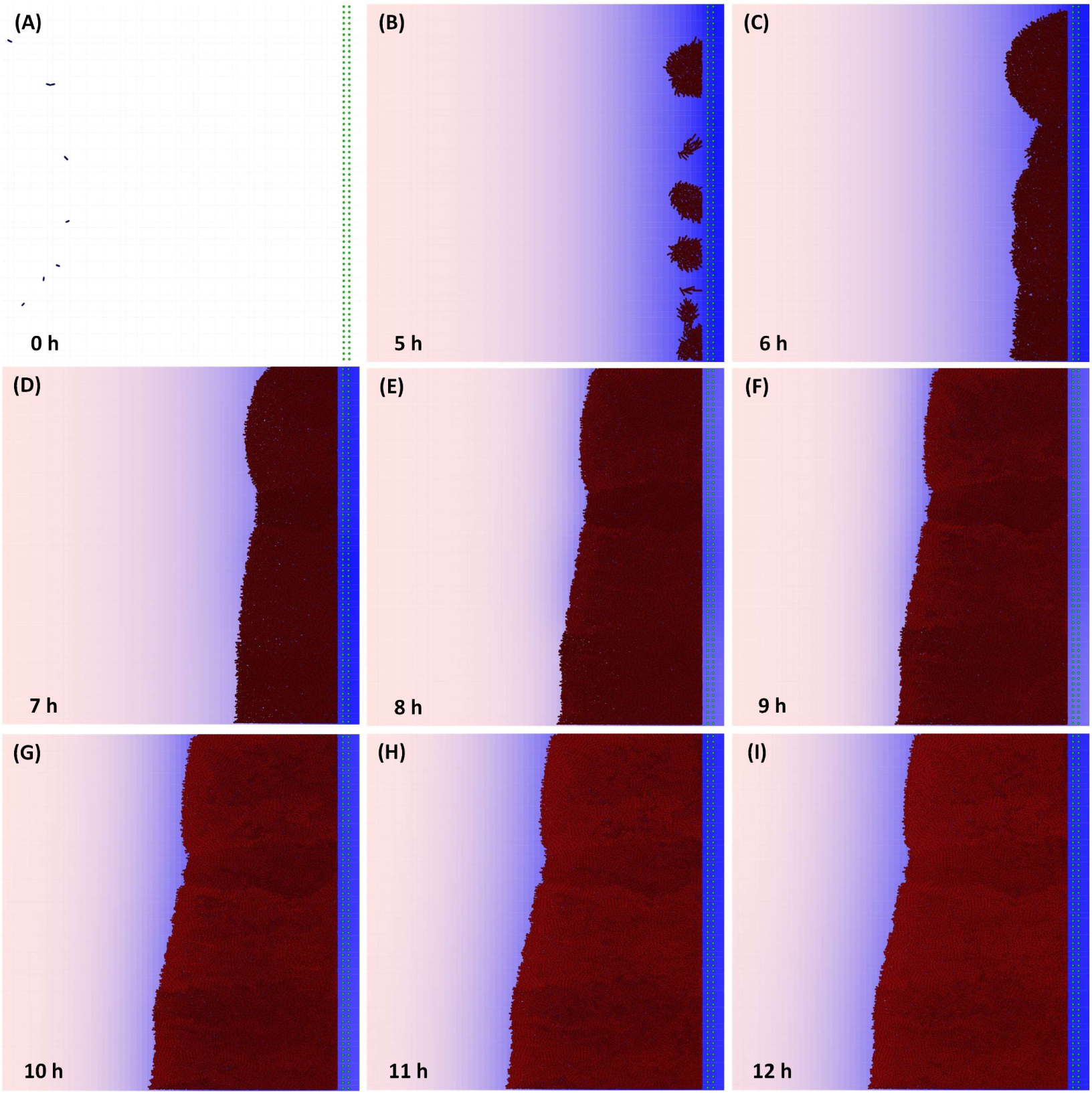
Elliptical cell arrangement with uniform NFs distribution (setup 2).(A) Initial setup of the simulation. At the left side of the cell surface, 8 motile cells are arranged elliptically. Motile cells are able to sense AI_2_ gradient produced by NFs. In the right side of the cell surface are the docked NFs (green circular objects). 128 NFs are distributed uniformly in two layers. During the simulation docked NFs (right side of each panel) produce AI_2_ which in turn is diffused onto the cell surface, blue color. (B)-(I) Motile cells reach the NFs site-the source of the signal-stop moving and start to grow and express their phenotype from 3 to 12 hours. See in the S2 movie the whole simulation.

The efficiency of the system is proportional to the HER2 density. As experimentally proved in [8], the efficiency of a similar system built in E. coli is proportional to the NF population. Our model complies with state-of-the-art knowledge. In Fig 11, we can observe that the number of cells expressing C increases to the number of NFs docked to the HER2. In S9 Fig, we present the results of a similar simulation setup but with different arrangement and number of NFs. It is obvious that by altering the NF distribution, we can obtain different efficiency in our system; however, the efficiency follows a logarithmic trend, see S10 Fig.

**Fig 11.**
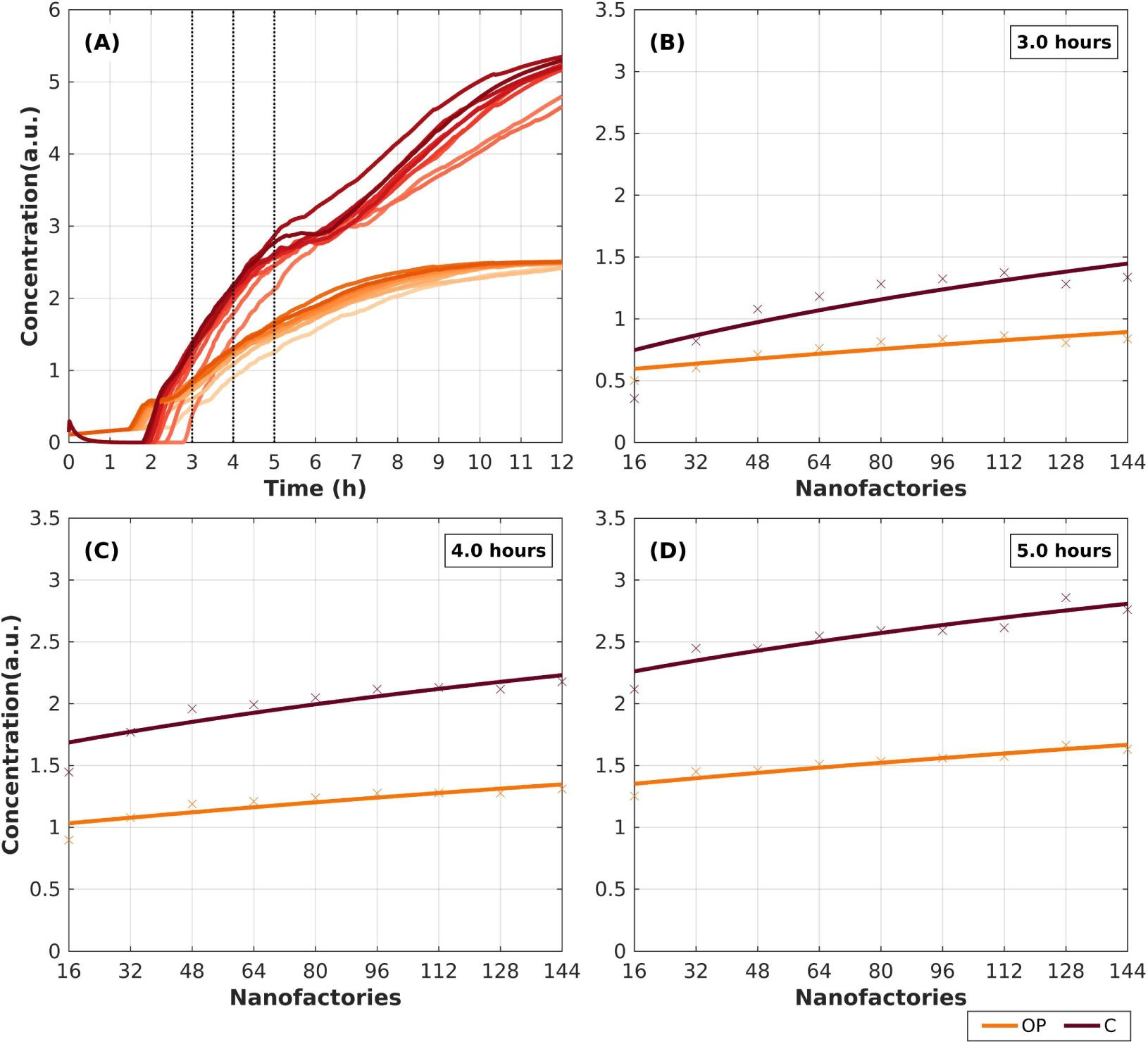
Expression of OP and C species with different number of NFs in setup 2. Colors represent the different species. (A) Evolution of OP and C expression in microbial community per NF setup. Light to dark orange represents the evolution of OP from 16 to 144 NFs. Light Red to maroon represents the evolution of C from 16 to 144 NFs. (B)-(D) The expression of OP and C per NFs setup at three time points (3, 4 and 5 hours, dashed lines in (A)). It is obvious that as the number of NFs increases the efficiency of the system increases logarithmically before the system reaches its steady state.

The spatial distribution except for the NFs population can play a significant role on the microcolony shape and consequently in virulent phenotype of the population. Thus, not only the population of NFs but also the spatial distribution of them must be considered in wet-lab experiments. The efficiency of such system to kill cancerous cells may depend on both NFs population and spatial distribution. In silico experimentation enables testing such hypothesis and designing better wet-lab experiments.

The ability to survey local surfaces and initiate gene expression based on feature density represents a new area-based switch in synthetic biology that it is believed to find use beyond the cancer model proposed herein. The assembled system comprising motile and sensing/actuating bacteria that (i) detect signal molecules emanating from receptors on tumor cells, (ii) migrate to the tumor cells, and (iii) initiate gene expression based on receptor density. The presented signal generation and cell recruiting design provides a tractable means to ensure site-specific gene initiation, providing a focused and predicted phenotype. Here, bacteria that naturally colonize specific niches, that favor virulence expression, are rewired (in silico) to perform switching functions based on prevailing feature densities or localized cell number (via self-generated AI_2_). We have intentionally designed the system as is, so that the incorporation of the control functions and output phenotype rely on minimally altered native circuits. The proposed case-study can be utilized in order to simulate in silico the state-of-the-art technology in experimental studies of human gut, named microfluidic Intestine Chip Models, proposed in [65,66].

## Methods

### Modeling methodology

To build a model that could be simulated (executable model), we first define the model reactions and the kinetic rate laws governing them. For the model to converge to the observed experimental behavior of the system, the underlying parameter values of the kinetic laws had to be precisely predicted. Prior to that, the starting state of the parameter search space for these values had to be defined. The latter step consisted of an extensive search in literature while the former step was implemented using stochastic global optimization algorithms. A depiction of the model’s mathematical definition and calibration steps is provided in Fig 12. In the Kinetic model development sub-section, we describe thoroughly the two first steps of the procedure, while in Methods section, we describe the rest of the procedure’s steps.

**Fig 12.**
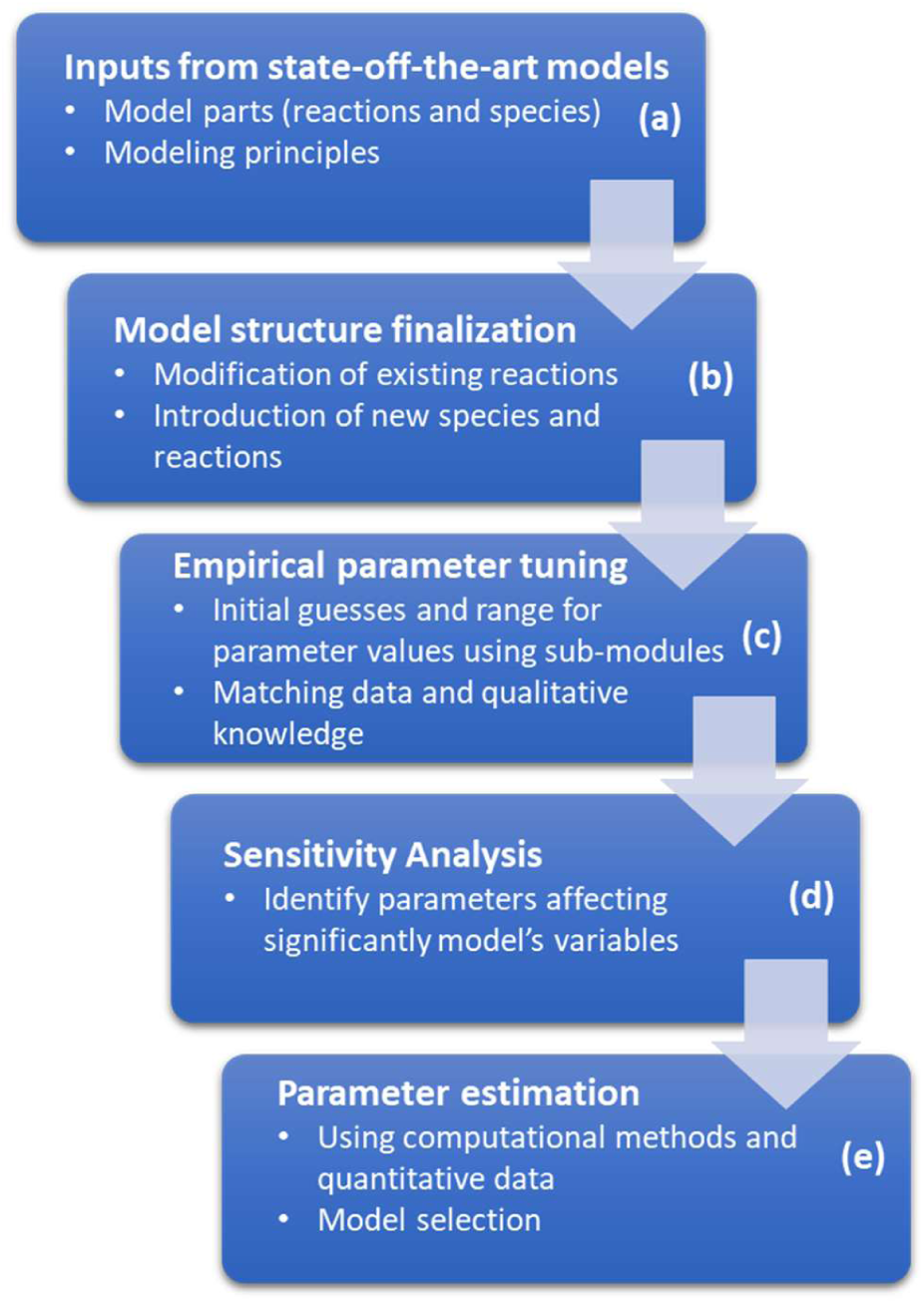
Modeling methodology. Model development (a)-(b) and calibration (c)-(e) procedure.

### Kinetic model

#### Modeling and Simulation

Initially, the model structure was captured in CellDesigner [67] to exploit the convenience of model definition in a graphical environment and the strength of Systems Biology Graphical Notation SBGN [68]. Then, the finalized model was imported (using the Systems Biology Workbench (SBW) [69]) to *COPASI* [54] (a publically available software) to be mathematically defined (definition of rate laws and assignment of parameter values for reactions and initial amounts/particle numbers for species). Furthermore, we used *COPASI* to compute a deterministic solution of the system of ordinary differential equations (presented in SM1 Table S1). Specifically, *COPASI* determines time course by using the LSODA integrator [55] which numerically evaluates a solution of the system. The parameters of the LSODA algorithm were chosen to have the default values set by the creators of the tool.

#### Empirical-preparatory tuning methodology

The developed models consists of 33 (30 in kinetic model) reactions which involve 14 (13 in kinetic model) distinct biochemical species. To decrease the complexity of the model we used experimental knowledge (whenever available) to group similar reactions and assign the same rate constants, leading to a total number of 45 free kinetic parameters. We set some parameter values directly from the literature, see S2 and S3 Tables. The rest of the parameters that were submitted for tuning in this step was decided based on empirical knowledge about their impact on the biological system’s behavior and on the observed initial simulation results too. The scope of this procedure was to manually tune a subset of the parameters to reach values that would drive the model to adequately converge to the qualitative knowledge for the biological system’s behavior, based on literature experimental findings. These parameter values were used as initial guesses for the final parameter estimation step using stochastic global optimization algorithms. The simulation algorithm used in this phase was deterministic (LSODA in *COPASI*).

The model simulation outputs were reviewed based on the following qualitative criteria: (a) AI-2 molecules present a sudden drop between 6 and 7 hours of growth following S. Typhimurium growth pattern (i.e. exponential growth). (b) HilA must be activated in order to achieve virulence into the simulations (c) LsrR represses InvF when no AI-2 is included into the environment. At each step of the empirical-preparatory tuning procedure of a parameter its value was adjusted following a trial and error procedure. The parameter values were first checked with respect to the Lsr Operon levels. Subsequently, for the values resulting into the expected Lsr Operon levels, it was attempted to reproduce the desired levels of virulence proteins, without affecting the previously matched levels of Lsr Operon. At the last step of this procedure, the parameter values were adjusted, having in mind the behavior of the model in the multicellular simulations too.

#### Sensitivity analysis

With respect to the model parameters, the sensitivity of the simulation results was systematically analyzed to determine if the model was sufficiently robust and able to capture the true dynamic behavior of the biological system. Moreover, the results of sensitivity analysis, after empirically tuning the model, were used to identify the set of the most significant parameters that would be re-estimated using global optimization algorithms. This resulted in the reduction of the computational cost of the procedure. Setting all parameters and initial conditions as variables, we calculated the scaled sensitivities with respect to the total Concentration of AI-2, Lsr Operon, SicA, SipC and in the cytoplasm (e.g., all species for which quantitative experimental data was available). This was done by numerical differentiation using finite differences, a tool integrated in *COPASI*. Scaled sensitivities were used since the previously mentioned species showed significant differences in their particle number levels. One should take note of the deterministic nature of the selected method and the fact that it only corresponds to local sensitivity analysis. This limitation was overcome by having obtained good estimates of parameter values prior to this step, derived by the empirical-preparatory manual tuning procedure. The sensitivity analysis results are summarized in S4 Table.

#### Calibration and Parameter estimation

We implement the parameter estimation task, using the experimental data (time-course experiments) found in [37,46,53,70,71], see S5 Table for more details. Based on the dynamics of the simulated regulatory networκ, we determined their initial values and their ranges, defining the search space. We defined these tolerance intervals based on our empirical knowledge on the suitability of the starting state. Furthermore, in order to reduce computational cost and simultaneously expedite the search of an optimal solution to our problem, those intervals had to be constrained. This was done through an iterative procedure where each range was adjusted accordingly after a series of search runs.

The next step of the calibration procedure was the estimation of the 38 parameters which we selected, using experimental data. The most frequently used approaches to search the parameter space are thoroguhly described in [72]. Here, we used Stochastic Global Optimization algorithms to estimate the selected parameters. Specifically, we chose Evolution Strategy [73], Particle Swarm [74] and Scatter Search [75] and Genetic Algorithm [76]. Taking into account the stochasticity which characterizes these algorithms, we conducted 200 parameter estimations with each algorithm. For this purpose we used the command line version of *COPASI*. After this step, we selected the best 100 fits (according to Root mean square error (RMSE) between the simulation results of each model, and the experimental data). Then, we calculated the mean and median values from the 100 estimations for each parameter and for each global optimization techniques used. Consequently, we created 8 models (one derived from the mean and one from the median of 100 parameter estimations for each algorithm), which we then evaluated. Finally, we choose the model with the lowest RMSE, i.e. the model with median value for each parameter from the Genetic Algorithm. The overview of the results appears in S6 Table.

### Individual-based model

To integrate the kinetic model into a multicellular framework following individual-based modeling approach [39,40] similar to the proposed in [32,34], we used *CellModeller* [56]. *CellModeller* includes models of biophysics, genetics, and intercellular signaling. Starting from single-cells it can simulate the development of colonies containing thousands of individual cells.

#### Cell Biophysics

Rod-shaped bacteria maintain highly consistent forms of roughly constant radius, with growth occurring exclusively on the long axis. This shape can be approximated by a cylinder capped with hemispherical ends, called a capsule. Under typical growth conditions, cells exhibit very little deformation so that they are well-approximated by rigid, elongating capsules. Then, we followed the formulated constrained rigid-body dynamics method, in which cell length is included as a degree of freedom, as proposed in [56]. Cells in [56] grow by an impulse adding length at each timestep, which is the only way energy is added to the system. The gamma parameter is the frictional drag on this growth. In our simulations we set gamma equal to 10, which is the default value proposed by [56]. A further consequence of gamma is how quickly cells stop growing due to physical forces. In our simulations, cells expend an amount of energy on growth at each timestep, but, this may need to go into pushing other cells out of the way. If there are enough cells blocking it, the cell will not be able to push them out of the way to grow. At low gamma values, cells are stopped from growing at low physical forces, and as gamma goes to infinity, cells get increasingly insensitive to physical forces and grow at the same rate in all situations.

#### Growth

The cells grow exponentially along the symmetry axis according to,

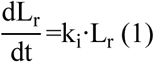

where *i* is the generation that the cell belongs to. As the cells grow the intracellular molecular concentrations decrease because of dilution. This dilution corresponds to the ratio of previous volume/current volume of a cell and it is applied to the species of each cell in each growth step, introducing noise in single-cell gene expression, as expected in such wet-lab experiments [41]. Once a cell reaches a certain threshold length (division length), it divides into two cells of almost identical size. We compute the division length when a new cell is generated (entering the simulation). The division length depends on its growth rate *k_i_* and division time *τ_i_*. Specifically, we consider a single-cell length model of the form *l*(*t*) = *l*(0)·e^kt^, as presented in [77,78], to estimate the length of a cell at a given time point. Given that we know the cell’s birth length *l*(0) and its growth rate *k* (i.e. cell elongation rate), then we compute the length at division *l*(τ), *l*(τ) = *l*(0)·e^kτ^.

In order to introduce stochasticity (noise) [42,43] on how the *S.* Typhimurium. single-cells grow in the simulation, we implemented and introduced a new divide function [56] to our model. Specifically, we generate randomly growth rate *k_i_* and division time *τ_i_* for each cell when it is inserted to the simulation, i.e. when it is generated by a division event, depending on its generation. Furthermore, at cell division we introduce randomness in order to break the axial symmetry of the system, to produce two daughter cells with slightly different sizes and imperfect alignment, as in [32].

#### Personalized kinetics extraction

To enable the proposed Individual based Modeling approach, we used *BaSCA* framework [79,80] so as to segment and track single-cells in S. Typhimurium time-lapse movies [42] and to reconstruct their lineage [81]. Given the extracted lineage tree of a colony and the estimated length of every cell at every time instant, we fit to each cell’s length trajectory (time series) an exponential model [77,78]. Specifically, we consider a single-cell length model of the form *l*(*t*) = *l*(0)·e^kt^, where *l*(0) is the birth length of a stalked cell and *k* is its growth rate (cell elongation rate). In this way we can capture the growth characteristics of each cell and estimate its *personalized* kinetic parameters.

Then, we extracted the best fit Gamma distributions of the personalized kinetic parameters, per cell generation (cell data from the different colonies are pooled). For each generation we fit a distribution out of Gamma, normal and lognormal and we selected the best according to BIC. Then according to the majority vote, we selected Gamma distribution because it was the best distribution for the majority of the generations. Generally, the gamma distribution is a flexible two-parameter distribution that belongs to exponential family and is used to model physical quantities that take positive values in microbiology, such as the cell division time (as in [43,82]]), the cell elongation rate (as in [43]) and cell division length (as in [78]). The estimated parameters are provided in S7 and S8 Tables. Single-cell analytics package [83] allowed us to quantify the stochasticity and examine inter and intra-generation variability in order to incorporate this knowledge to individual-based modeling approach.

Intracellular Dynamics and Signaling.

#### Adaptation of kinetic model equations

With few exceptions, the previously described ODEs were repurposed without modification in the Individual based model simulations. The exchange of AI-2 between the environment and a cell was localized to the space in which the cell center was found at the beginning of the time step. This allowed a synchronized update of the final grid concentration at the end of the time step. Furthermore, the AI-2 dilution/concentration factor was adjusted from 10^12^ to 64 to account for the difference between the milliliter volume associated with cell concentration and the implied volume of grid elements.

#### Intracellular dynamics

We use ordinary differential equations (ODEs) to approximate intracellular dynamics. Variability between cells are simulated in our model by introducing randomness at the partitioning of a molecular species and length between daughter cells during cell division and by introducing stochasticity at elongation rate and division time of each cell, as discussed previous section. Here, we formulated a system of differential equations, the latter describing the rate of change of a species. In general, and most commonly for genetic circuits, this system is nonlinear, where the rate of change of the vector of species 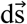 is of the form:

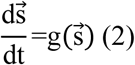

where the function *g* defines the rate of change of each species, see S1 Table, inside each cell.

#### Cell signaling

In our modelling approach, we used cell signaling to explore multicellular organization [84]. In microbial communities, cells communicate via quorum sensing ligands that diffuse through the environment’s medium, after being secreted from the cell. In our simulations, microbial communities grow into agar plate environment, where transport processes, such as advection and diffusion, usually occur. We include such signaling through the medium in our model with a general linear transport operator **T**, as proposed in [56]:

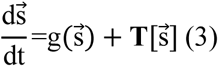

where 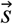 is now composed of some species that are within a cell (and can not be transported οutside of cell membrane) and those outside the cell, which are subject to the operator T. Specifically, for diffusion, the operator **T** ≡ **D**∇^2^, where **D** is a diagonal matrix of diffusion coefficients for external AI2 species, whereas for internal cell species the corresponding elements of **D** are zero.

As proposed in [56], we discretized this system on a regular 3-dimensional grid for species in the medium and separate variables representing cell autonomous species. Cell positions are interpolated linearly in the spatial grid, and each cell can sense its local AI-2 concentration (extracellularly). We solve the resulting system of nonlinear partial differential equations with the aim of a modified Crank-Nicholson method [57].

## Acknowledgements

The first author acknowledges the support of the Alexander S. Onassis Public Benefit Foundation (scholarship number GZJ030/2013-2014).

## Author Contributions

Conceived and designed the experiments: ADB AT ESM. Performed the experiments: ADB. Analyzed the data: ADB. Contributed reagents/materials/analysis tools: ADB. Wrote the paper: ADB ESM.

## Notes

### Competing Interest Statement

The authors have declared no competing interest.

