## Supplementary material for "Individual-Based Modeling of Microbial Communities Integrating Genetic Mechanisms: A Case Study of LuxS-Mediated Quorum Sensing in Salmonella Typhimurium": S1 Text

### Supporting Information

#### Model definition

##### Reactions

Following the notation, we used in the main text, we present the reactions and their kinetic laws of the kinetic and individual based models, see S1 Fig. The reactions in the green box represent the QS dynamics, while the reactions in the magenta box represent the TTSS-1 dynamics. The reactions in gray color represent the dynamics outside the cell's cytoplasm, thus they are included only in individual-based modelling approach, for more details refer to the Kinetic model development subsection (in the main text).

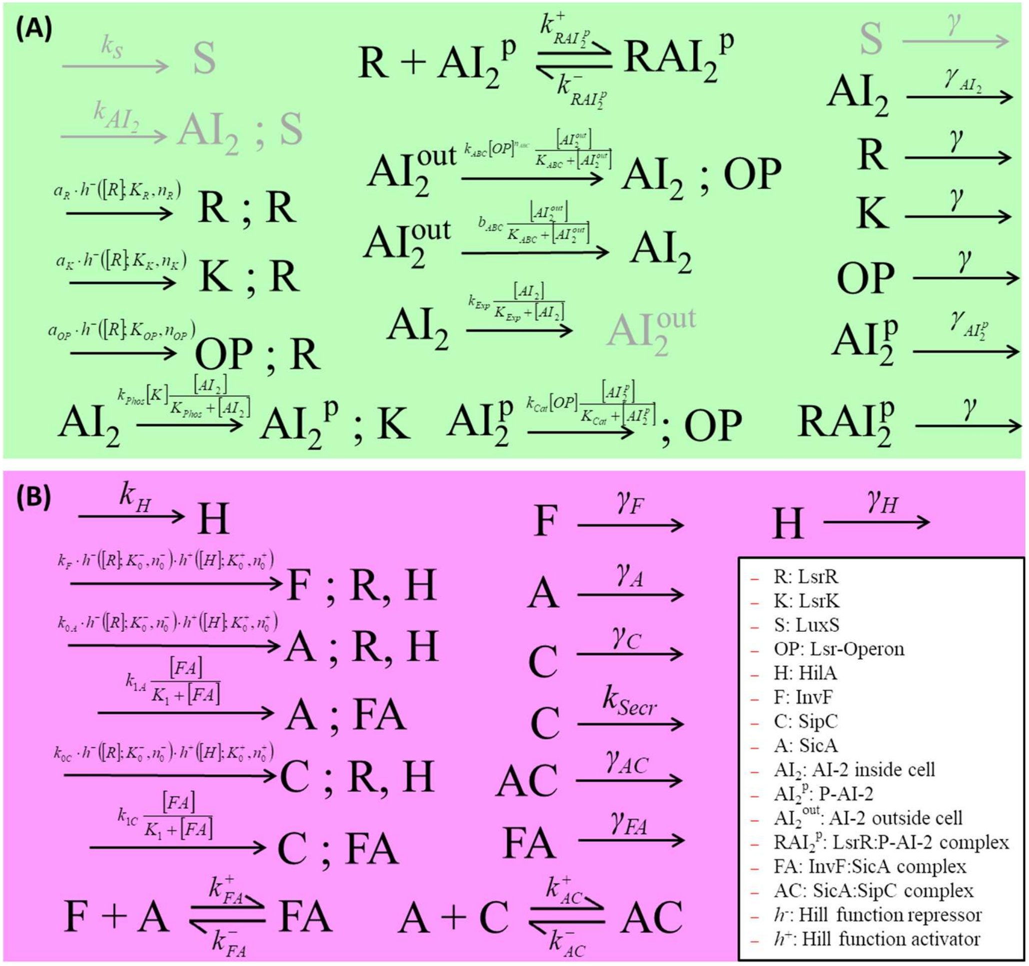


**S1 Fig. Set of reactions of the proposed models.** (A) Reactions and kinetic laws (above arrows) of the LuxS-mediated QS in *S.* Typhimurium. (B) Reactions and the kinetic laws of the TTSS-1 in *S.* Typhimurium. In the legend, we present the abbreviations of species names and the Hill function types found in our models. Notice that regulatory protein R (LsrR) exists in both panels because it regulates both the QS and TTSS-1 mechanisms. Reactions in gray color participate only in individual-based model approach.

#### Differential equations

Following the notation, we used in the main text, we present the system of ordinary differential equations of the kinetic and individual-based model in S1 Table.

**S1 Table. Ordinary differential equations of each modelling approach.**

| Model | Equation |
|---|---|
| **Single-cell and Multicellular** | $\frac{d[R]}{dt}=k_R\frac{1}{1+([R]/K_{RK})^{n_{RK}}}-\left(k^{+}_{RAI^p_2}[R][AI^p_2]-k^{-}_{RAI^p_2}[RAI^p_2]\right)-\gamma[R]$<br>$\frac{d[K]}{dt}=k_K\frac{1}{1+([R]/K_{RK})^{n_{RK}}}-\gamma[K]$<br>$\frac{d[OP]}{dt}=k_{OP}\frac{1}{1+([R]/K_{OP})^{n_{OP}}}-\gamma[OP]$<br>$\frac{d[AI^p_2]}{dt}=k_{Phos}[K]\frac{[AI_2]}{K_{Phos}+[AI_2]}-\left(k^{+}_{RAI^p_2}[R][AI^p_2]-k^{-}_{RAI^p_2}[RAI^p_2]\right)-k_{Cat}[OP]\frac{[AI^p_2]}{K_{Cat}+[AI^p_2]}-\gamma_{AI^p_2}[AI^p_2]$<br>$\frac{d[RAI^p_2]}{dt}=k^{+}_{RAI^p_2}[R][AI^p_2]-k^{-}_{RAI^p_2}[RAI^p_2]-\gamma[RAI^p_2]$<br>$\frac{d[H]}{dt}=k_H-\gamma[H]$<br>$\frac{d[F]}{dt}=k_F\frac{([H]/K^{+}_0)^{n^{+}_0}}{1+([H]/K^{+}_0)^{n^{+}_0}}\cdot\frac{1}{1+([R]/K^{-}_0)^{n^{-}_0}}-(k^{+}_{FA}[F][A]-k^{-}_{FA}[FA])-\gamma_F[F]$<br>$\frac{d[A]}{dt}=k_{0A}\frac{([H]/K^{+}_0)^{n^{+}_0}}{1+([H]/K^{+}_0)^{n^{+}_0}}\cdot\frac{1}{1+([R]/K^{-}_0)^{n^{-}_0}}+k_{1A}\frac{[FA]}{K_1+[FA]}-(k^{+}_{FA}[F][A]-k^{-}_{FA}[FA])-(k^{+}_{AC}[A][C]-k^{-}_{AC}[AC])-\gamma_A[A]$<br>$\frac{d[C]}{dt}=k_{0C}\frac{([H]/K^{+}_0)^{n^{+}_0}}{1+([H]/K^{+}_0)^{n^{+}_0}}\cdot\frac{1}{1+([R]/K^{-}_0)^{n^{-}_0}}+k_{1C}\frac{[FA]}{K_1+[FA]}-(k^{+}_{AC}[A][C]-k^{-}_{AC}[AC])-k_{Secr}[C]-\gamma_C[C]$<br>$\frac{d[FA]}{dt}=k^{+}_{FA}[F][A]-k^{-}_{FA}[FA]-\gamma_{FA}[FA]$<br>$\frac{d[AC]}{dt}=k^{+}_{AC}[A][C]-k^{-}_{AC}[AC]-\gamma_{AC}[AC]$ |
| **Single-cell** | $\frac{d[AI_2]}{dt}=k_{Phos}[K]\frac{[AI_2]}{K_{Phos}+[AI_2]}+k_{PTS}\frac{[AI^{out}_2]}{K_{PTS}+[AI^{out}_2]}+k_{ACBD}[OP]^{n_{ACBD}}\frac{[AI^{out}_2]}{K_{ACBD}+[AI^{out}_2]}-k_{Exp}\frac{[AI_2]}{K_{Exp}+[AI_2]}-\gamma_{AI_2}[AI_2]$<br>$[AI^{out}_2]=\begin{cases} a-\frac{a}{\lambda}\cdot\lvert t-t_{max}\rvert & \text{if } \frac{a}{\lambda}\cdot\lvert t-t_{max}\rvert < a \\ 0 & \text{else}\end{cases}$ |

| | |
|---|---|
| **Multicellular** | $\frac{d[S]}{dt}=k_S-\gamma[S]$ <br> $\frac{d[AI_2]}{dt}=k_{AI_2}[S]-k_{Phos}[K]\frac{[AI_2]}{K_{Phos}+[AI_2]}+$ <br> $+\left(k_{PTS}\frac{[AI_2^{out}]}{K_{PTS}+[AI_2^{out}]}+k_{ACBD}[OP]^{n_{ACBD}}\frac{[AI_2^{out}]}{K_{ACBD}+[AI_2^{out}]}-k_{Exp}\frac{[AI_2]}{K_{Exp}+[AI_2]}\right)\frac{S_{cell}}{V_{cell}}-\gamma_{AI_2}[AI_2]$ <br> $\frac{d[AI_2^{out}]}{dt}=\left(-k_{PTS}\frac{[AI_2^{out}]}{K_{PTS}+[AI_2^{out}]}-k_{ACBD}[OP]^{n_{ACBD}}\frac{[AI_2^{out}]}{K_{ACBD}+[AI_2^{out}]}+k_{Exp}\frac{[AI_2]}{K_{Exp}+[AI_2]}\right)\frac{S_{cell}}{V_{grid}}-D\nabla^2[AI_2^{out}]$ |

Species concentrations are denoted with [ ], e.g. concentration of species S is denoted as [S]. Rate constants are presented in detail in S2 Table. $S_{cell}$: cell area, $V_{cell}$: cell volume, $V_{grid}$: grid volume and D: diffusion constant

Following the notation presented in the main text, we present the parameters' values of the kinetic and individual-based model. In S2 Table we present the parameters concerning the QS species while in S3 Table we present the parameters concerning TTSP-1 species. The majority of the model's parameters were estimated using experimental data found in literature, see in Calibration and Parameter estimation subsection in the main text. Notice that in S2 and S3 Table description columns, the parameters that were set from literature include the corresponding reference.

**S2 Table. QS module rate constants.**

| Parameter | Description | Value |
|---|---|---|
| $^*k_S$ | LuxS synthesis rate | $0.030728$ a.u.$\cdot$min$^{-1}$ |
| $k_{OP}$ | Lsr-operon synthesis rate | $0.0505232$ a.u.$\cdot$min$^{-1}$ |
| $k_R$ | LsrR synthesis rate | $0.00688882$ a.u.$\cdot$min$^{-1}$ |
| $k_K$ | LsrK synthesis rate | $0.00688882$ a.u.$\cdot$min$^{-1}$ |
| $^*k_{AI_2}$ | AI-2 synthesis rate | $0.2365253073$ min$^{-1}$ |
| $k_{Phos}$ | AI-2 phosphorylation rate | $12.64735$ min$^{-1}$ |
| $k_{Cat}$ | P-AI-2 catalysis rate | $16.81485$ min$^{-1}$ |
| $k^{+}_{RAI_2^P}$ | P-AI-2/LsrR binding rate | $30.27815$ a.u.$^{-1}\cdot$min$^{-1}$ |
| $k^{-}_{RAI_2^P}$ | P-AI-2/LsrR disassociation rate | $0.00243164$ min$^{-1}$ |
| $n_{OP}$ | Cooperativity coefficient of Lsr operon | 5.12211 |
| $n_{RK}$ | Cooperativity coefficient of LsrRK operon | 5.12211 |

| | | |
|---|---|---|
| $n_{ACBD}$ | Cooperativity coefficient of LsrACBD | 4.529005 |
| $k_{PTS}$ | AI-2 import rate by PTS mechanism | 781.4665 $min^{-1}$ |
| $k_{ACBD}$ | AI-2 import rate by LsrACBD | 5.670385 $a.u.^{-1} \cdot min^{-1}$ |
| $k_{Exp}$ | AI-2 export rate | 3.54443 $a.u. \cdot min^{-1}$ |
| $K_{OP}$ | Dissociation coefficient of Lsr operon | 0.2210345 a.u. |
| $K_{RK}$ | Dissociation coefficient of LsrRK operon | 18.5479 a.u. |
| $K_{Phos}$ | Dissociation constant of AI-2 phosphorylation | 578.437 a.u. |
| $K_{Cat}$ | Dissociation constant of P-AI-2 catalysis | 36.98545 a.u. |
| $K_{Exp}$ | Dissociation constant of AI-2 exportation | 592.774 a.u. |
| $K_{ACBD}$ | Dissociation constant of AI-2 importation by LsrABC | 96.66255 a.u. |
| $K_{PTS}$ | Dissociation constant of AI-2 importation by PTS mechanism | 57.17795 a.u. |
| a** | $AI_2^{out}$ maximal concentration [S1] | 69 x $10^3$ |
| $t_{max}$** | timepoint when $AI_2^{out}$ reaches maximal concentration [S1] | 240 min |
| $\gamma$ | degradation/dilution [S2] | 0.02 $min^{-1}$ |
| $\gamma_{AI_2^p}$ | P-AI-2 degradation rate [S3] | 0.3 $min^{-1}$ |
| $\gamma_{AI_2}$ | AI-2 degradation rate [S3] | 0.15 $min^{-1}$ |

*parameters used only in individual-based model approach

**parameters used only in kinetic model approach.

a.u. = arbitrary concentration units.

#### S3 Table. Virulence module rate constants.

| Parameter | Description | Value |
|---|---|---|
| $k_H$ | HilA synthesis rate | 0.0463577 $a.u. \cdot min^{-1}$ |
| $k_{0A}$ | SicA external promoter synthesis rate | 2.63596 $a.u. \cdot min^{-1}$ |
| $k_{0C}$ | SipC external promoter synthesis rate | 6.28015 $a.u. \cdot min^{-1}$ |
| $k_F$ | InvF synthesis rate | 2.061625 $a.u. \cdot min^{-1}$ |
| $k_C$ | SipC synthesis rate | 61.9536 $a.u. \cdot min^{-1}$ |
| $k_A$ | SicA synthesis rate | 33.44115 $a.u. \cdot min^{-1}$ |
| $k_{FA}^{+}$ | InvF:SicA forward rate | 0.01340425 $a.u.^{-1} \cdot min^{-1}$ |

| | | |
|---|---|---|
| $k^{-}_{FA}$ | InvF:SicA backward rate [S4] | 13.7225 $min^{-1}$ |
| $k^{+}_{AC}$ | Sic:SipC forward rate | 0.283012 $a.u.^{-1}\cdot min^{-1}$ |
| $k^{-}_{AC}$ | SicA/SipC backward rate [S5] | 4.77204 $min^{-1}$ |
| $n^{R}_{0}$ | Cooperativity coefficient of inv operon (external promoter) by LsrR | 5.15456 |
| $n^{H}_{0}$ | Cooperativity coefficient of inv operon (external promoter) by HilA | 5.15456 |
| $K^{H}_{0}$ | Dissociation coefficient of inv operon (external promoter) by HilA | 11.81735 a.u. |
| $K^{R}_{0}$ | Dissociation coefficient of inv operon (external promoter) by LsrR | 0.3120015 a.u. |
| K | Dissociation constant inv operon (internal promoter) | 3.921615 a.u. |
| $k_{Secr}$ | Secretion rate of SipC [S6][S7] | 0.1 $min^{-1}$ |
| $\gamma_H$ | HilA degradation/dilution [S8] | 0.067 $min^{-1}$ |
| $\gamma_F$ | InvF degradation/dilution [S9] | 0.022 $min^{-1}$ |
| $\gamma_C$ | SipC degradation/dilution [S10][S11] | 0.139 $min^{-1}$ |
| $\gamma_A$ | SicA degradation/dilution [S4] | 0.046 $min^{-1}$ |
| $\gamma_{FA}$ | InvF/SicA degradation/dilution [S9] | 0.022 $min^{-1}$ |
| $\gamma_{AC}$ | SicA/SipC degradation/dilution [S9] | 0.022 $min^{-1}$ |

a.u. = arbitrary concentration units.

#### Sensitivity analysis

For this procedure, we used the already implemented software that is offered by COPASI [S12]. Our target – system‘s output was the value of OP (Lsr operon), F (InvF), A (SicA), C (SipC) (these were the species we had experimental data of). We set all of the system‘s parameters to be considered sensitive, thus every parameter was changed in order for us to see the effect it had in the system‘s output. We have to note here that the method offered by COPASI, refers to local sensitivity analysis. The results we got from this procedure came in two flavors, scaled and unscaled. We decided to proceed with the scaled results; We have to note here that in sensitivity analysis the scaled values of the parameters can be signed. That means that a parameter might be the most sensitive but have a negative value. The scaled results are shown in the following Table.

**S4 Table. Sensitivity Analysis Results**

| | **Sensitivity values (scaled sensitivities)** |
|---|---|

| **Parameters/ Species Initial Concentration** | **Criterion I: [OP]** | **Criterion II: [F]** | **Criterion III: [A]** | **Criterion IV: [C]** |
|---|---|---|---|---|
| $\gamma$ | 2.499680 | -0.004477 | 0.038872 | 0.055529 |
| $\gamma_C$ | -0.000040 | -0.000005 | 0.104451 | -0.708970 |
| $\gamma_A$ | -0.000028 | 0.000162 | -15.199300 | -14.823500 |
| $\gamma_{AI_2}$ | -0.117947 | -0.030822 | -0.062352 | -0.091334 |
| $\gamma_{AI_2^p}$ | -0.007068 | -0.002140 | -0.002849 | -0.003690 |
| $\gamma_F$ | -0.000007 | -7.070270 | -5.586100 | -12.395200 |
| $\gamma_{FA}$ | -0.000003 | -0.022025 | -0.067023 | -0.089772 |
| $\gamma_H$ | -0.000031 | -1.405990 | -5.181440 | -6.678550 |
| $\gamma_{AC}$ | -0.000034 | -0.000006 | -0.488513 | -0.483956 |
| $k_{Secr}$ | -0.000005 | 0.000048 | 0.059440 | -0.523265 |
| a | 0.108803 | 0.028906 | 0.063508 | 0.090104 |
| $K_0^H$ | -0.000003 | -0.954213 | -4.412280 | -5.331910 |
| $K_0^R$ | -0.000046 | 0.070187 | 0.186178 | 0.278479 |
| K | 0.000005 | -0.000044 | -3.892270 | -4.853150 |
| $K_{OP}$ | 4.340520 | 0.000948 | 0.000303 | 0.000550 |
| $K_{RK}$ | -0.000001 | -0.000009 | -0.000090 | 0.000134 |
| $K_{ACBD}$ | -0.016065 | -0.005895 | -0.011029 | -0.018817 |
| $K_{Cat}$ | 0.022685 | 0.008304 | 0.015940 | 0.023789 |
| $K_{Exp}$ | 0.004693 | 0.001224 | 0.003068 | 0.004474 |
| $K_{Phos}$ | -0.102872 | -0.027375 | -0.059666 | -0.086607 |
| $K_{PTS}$ | -0.092711 | -0.023073 | -0.052039 | -0.073396 |
| $k_{0A}$ | -0.000004 | -0.000063 | 0.817730 | 0.811678 |
| $k_{0F}$ | -0.000032 | 1.000060 | 3.910000 | 4.879530 |
| $k^-_{RAI_2^p}$ | -0.006485 | -0.001707 | -0.002584 | -0.005644 |
| $k^+_{RAI_2^p}$ | 0.029034 | 0.010256 | 0.021665 | 0.032404 |
| $k_A$ | -0.000043 | -0.000006 | 4.074980 | 4.046170 |
| $k^+_{FA}$ | -0.000031 | 0.000046 | 0.002102 | -0.001372 |
| n | -0.000001 | 0.033161 | 0.065042 | 0.096400 |
| $n_{OP}$ | -1.770930 | 0.000416 | 0.002077 | 0.003907 |
| $n_{ACBD}$ | 0.064075 | 0.024245 | 0.045672 | 0.066623 |
| $k_{0C}$ | -0.000028 | -0.000008 | -0.054072 | -0.048894 |

| | | | | |
|---|---|---|---|---|
| $k_{AC}^{+}$ | -0.000006 | 0.000031 | -0.001030 | -0.001701 |
| $k_C$ | -0.000021 | -0.000058 | -0.124131 | 0.866969 |
| $k_H$ | 0.000007 | 0.955138 | 4.434220 | 5.362550 |
| $[R]_0$ | -0.000077 | -0.000402 | -0.014127 | -0.013624 |
| $[AI_2]_0$ | -0.000001 | -0.000020 | 0.000503 | 0.001610 |
| $[H]_0$ | -0.000027 | 0.000018 | 0.000748 | 0.002114 |
| $[F]_0$ | -0.000001 | 0.000015 | 0.000527 | -0.000250 |
| $[K]_0$ | 0.000006 | -0.000002 | 0.000266 | -0.000147 |
| $[OP]_0$ | 0.000000 | -0.000007 | 0.000019 | -0.000267 |
| $[RAI_2^p]_0$ | -0.000004 | -0.000056 | -0.000958 | -0.000053 |
| $[AI_2^p]_0$ | 0.000005 | 0.000031 | 0.002097 | 0.002503 |
| $[A]_0$ | -0.000002 | -0.000018 | 0.000206 | 0.002285 |
| $[FA]_0$ | -0.000041 | -0.000007 | 0.000891 | 0.001749 |
| $[AC]_0$ | -0.000030 | 0.000001 | 0.000028 | 0.000110 |
| $[C]_0$ | -0.000028 | 0.000036 | 0.001171 | 0.001856 |

### Results

#### Model validation

We did knock-outs of luxS and lsr genes in silico and we observed how they affect lsr operon expression in individual-based modelling approach.

First, we simulated knock-outs experiments of *luxS* and lsr genes in the kinetic model and we monitor how they affect lsr operon expression, see S2 Fig. Simulation time in all the experiments is 12 hours.

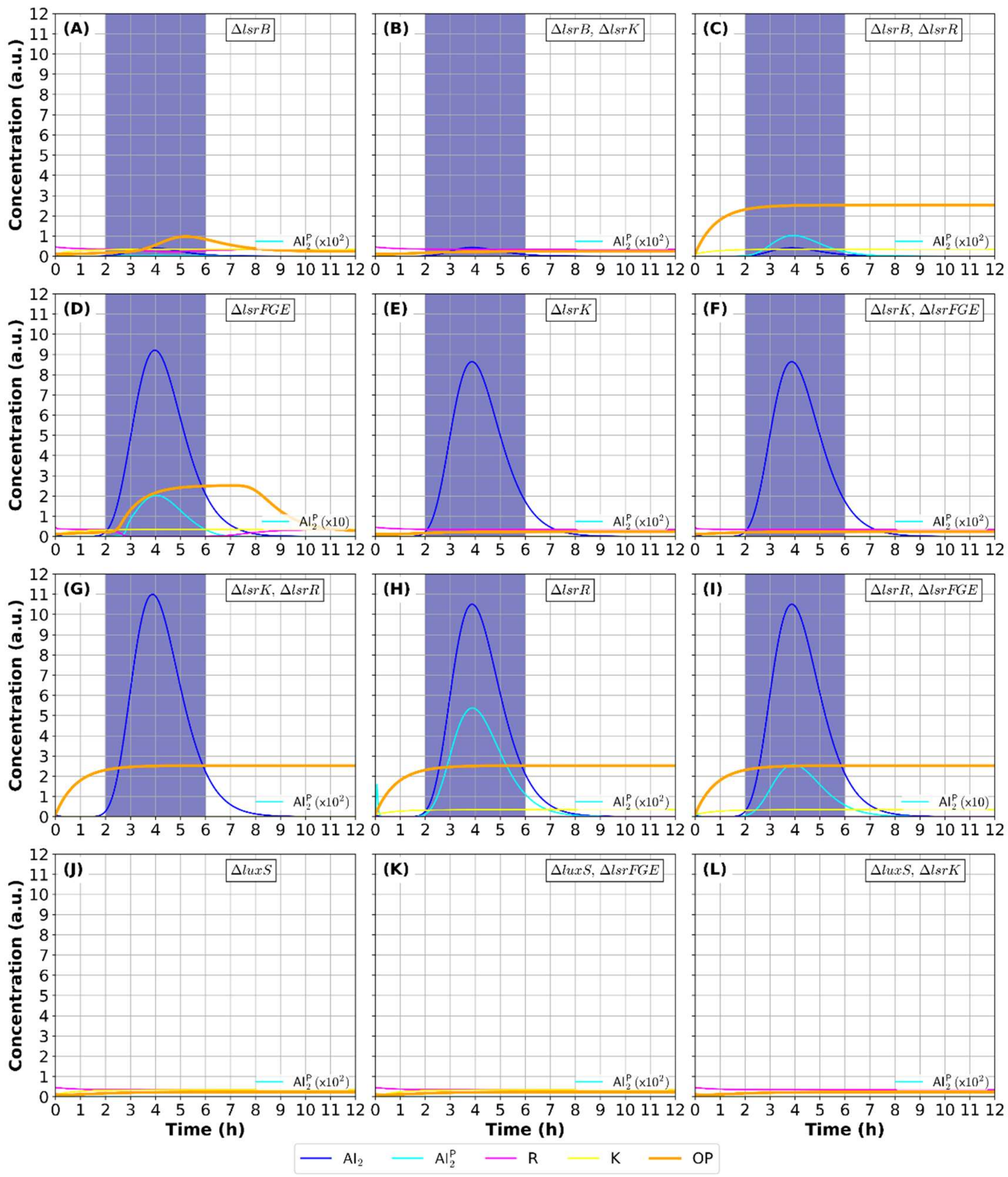

**S2 Fig. lsr genes single and double knock-outs effect on lsr operon species expression.** Colors represent the different species. Dark blue region represents the input $AI_2^{out}$ pulse. Time-course results for 12 hours experimental time. In each knock-out in-silico experiment the OP (orange curve) fold of expression (at steady-state) matches qualitatively with the corresponding knock-out wet-lab experiments presented in *Taga et al.* **[46]**.See text for details.

**Δ*lsrB*** [46] **/ Δ*lsrB* and Δ*lsrK*** [46]**.** In S2 Fig (A) and (B), we can observe that OP expression follows similar trend. In the former case there is very low $AI_2^p$ introduced to the system (only by AI-2 flux) because there is no LsrB to import it through the transmembrane protein complex, thus we obtain very low OP expression. In the latter case, there is no OP production because there is no K produced to phosphorylate $AI_2$, to relieve lsr operon repression by R. Both our findings concur with the experiments presented in [46] (Fig 5).

**Δ*lsrB* and Δ*lsrR*** [46]. In S2 Fig (C), we can observe that OP is produced from the beginning of the in silico experiment, despite the very low concentration of $AI_2$ and thus $AI_2^p$ in the system due to LsrB absence. Obviously, this happens because there is no R to repress OP production. Our findings on OP expression are in agreement - from a qualitative viewpoint - with the experimental findings presented in [46] (Fig 5).

**Δ*lsrFGE*** [58,46]. In S2 Fig (D), we can see that $AI_2^p$ is overexpressed because there is no LsrFGE to degrade it; thus, R binds to it massively, loosening the repression of lsr operon, resulting in OP expression increase (slightly in folds of expression), as shown in [46] (Fig 5). Furthermore, OP expression follows very similar trend (time-course), as it has been shown in Marques *et al*. [58] (Fig 2 (A) in main text).

**Δ*lsrK*** [58,46] **/ Δ*lsrK* and Δ*lsrFGE*** [46]. In S2 Fig (E) and (F) we can observe that $AI_2^p$ cannot be formed since there is no K to phosphorylate $AI_2$. There is no $RAI_2^p$ complex production. Thus, R represses OP and itself too. It is obvious that ΔlsrFGE makes no difference to the simulation dynamics. In both experiments there is no $AI_2^p$ produced so there is nothing to be degraded by LsrFGE. Consequently, in both experiments we have no expression of OP, as it has been shown experimentally in [46] (Fig 5). Furthermore, OP expression follows a very similar trend (time-course), as it has been shown in Marques *et al*. [58] (Fig 2 (A) in main text).

**Δ*lsrR*** [46] **/ Δ*lsrR* and Δ*lsrK*** [46]**.** In lsrR single knock-out, we obtain early production of OP and $AI_2^p$ because R cannot be produced and thus represses the lsr operon transcription, see S2 Fig (H). In ΔlsrR and ΔlsrK, we get the same result as regards OP expression, but there is no $AI_2$ phosphorylation due to K absence, see S2 Fig (G). Our findings on OP expression concur qualitatively with the experimental findings presented in [46] (Fig 5).

**Δ*lsrFGE* and Δ*lsrR*** [46]**.** Again, in the knock-out experiment, OP is highly expressed from the beginning of the experiment because there is no R repressing it. It is obvious that ΔlsrFGE makes no difference in OP dynamics when lsrR gene is deleted. However, because there is no $AI_2^p$ degradation, we can observe that $AI_2^p$ is overexpressed, S2 Fig (I). Our findings on OP expression agree qualitatively with the experimental findings presented in [46] (Fig 5).

**Δ*luxS*, Δ*luxS* and Δ*lsrFG*E, Δ*luxS* and Δ*lsrK*** [46]**.** In S2 Fig (J), we obtain very low OP expression, because R keeps repressing the model's species. There is no $AI_2$ production inside cell, consequently there is no $AI_2^{out}$ in the micro-environtment. The same results occur in double knock-out experiments ΔluxS and ΔlsrFGE, ΔluxS and ΔlsrK, see S2 Fig (K) and (L) respectively. Our results on R fold of expression are consistent with experimental findings in [46] Fig 1 and Fig 4 (A).

In S3 Fig, we present the corresponding results of the Ibm.

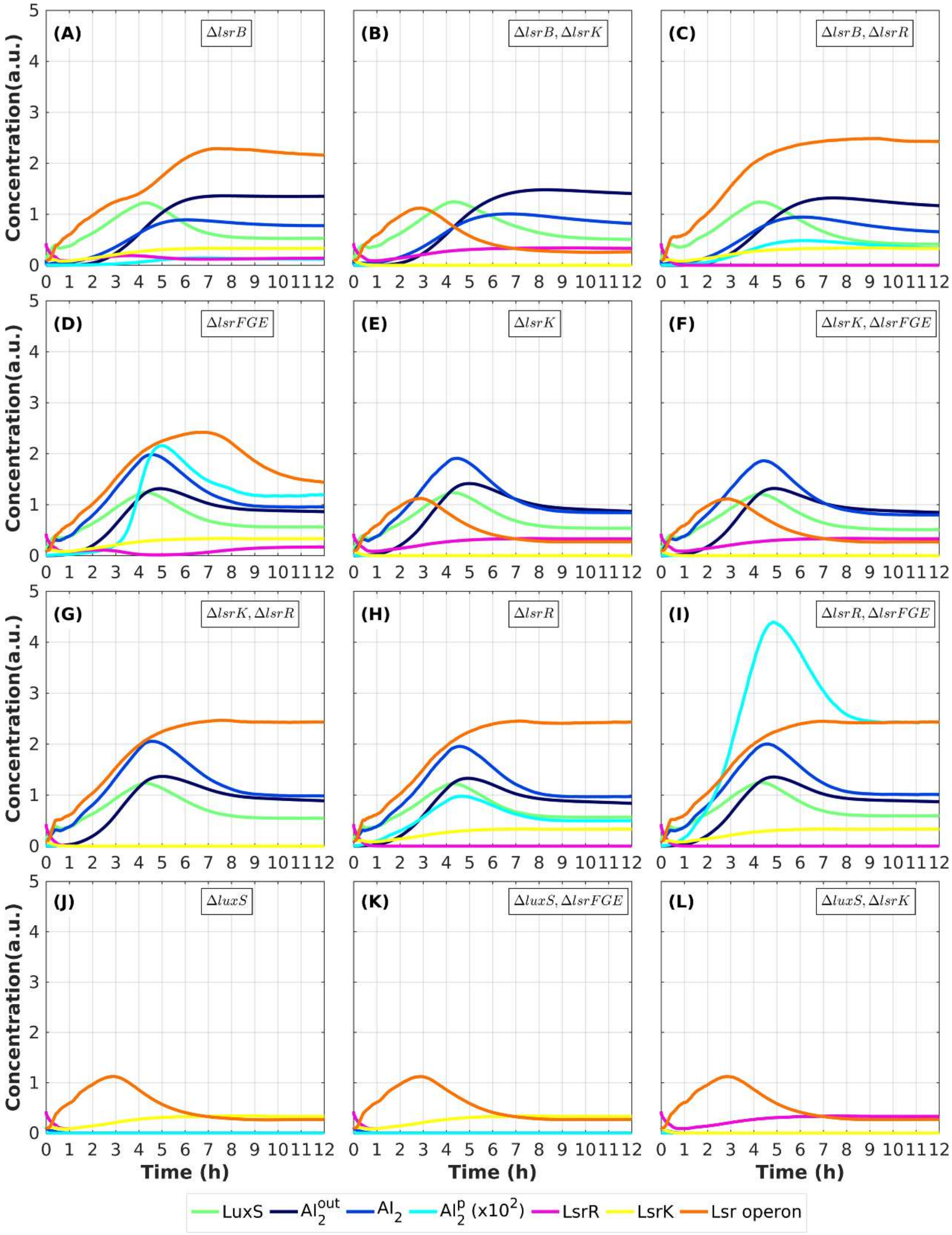


**S3 Fig. lsr genes single and double knock-outs effect on lsr operon species expression in Ibm.** Colors represent the different species. Time-course results for 12 hours experimental time. In each knock-out in-silico experiment the OP (orange curve) fold of expression (at steady-state) matches qualitatively with the corresponding knock-out wet-lab experiments presented in *Taga et al.* [S1]. See main text for details.

Then we proceed with knock-outs experiments of *luxS* and lsr genes in silico and we monitor how they affect TTSS-1 species expression, see S4 Fig for kinetic model results. Simulation time in all the experiments is 12 hours.

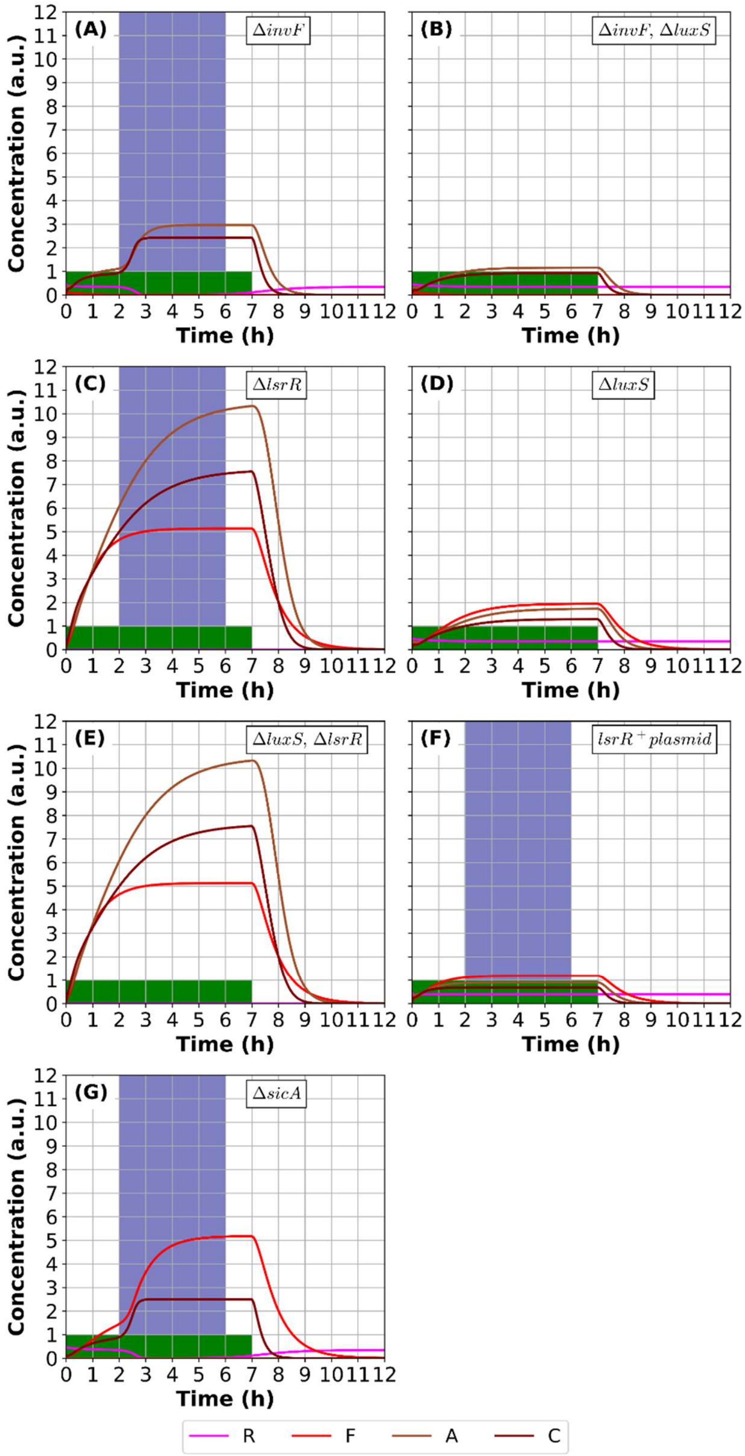


**S4 Fig. lsrR, luxS, invF and sicA knock-outs effect on virulence.** Colors represent the different species. Dark blue region represents the input $AI_2^{out}$ pulse. Green region represents the interval that H is activated. Time-course results for 12 hours experimental time. In each knock-out in silico experiment R and TTSS-1 species fold of expression (at steady-state) matches qualitatively with the corresponding knock-out wet-lab experiments presented in *Choi et al.* **[38]**. See text for details.

**Δ*invF*, Δ*luxS*, Δ*invF* and Δ*luxs* [38]**. When F is not expressed the downstream feedback loop is deactivated, thus TTSS-1 species expression is reduced. S controls virulence activity by regulating invF expression while R exists in the environtment. In Δ*invF*, Δ*luxS* or Δ*invF* and Δ*luxS*, A and C expression is reduced comparing to the WT expression, as shown in S4 Fig (A), (B) and (D)**.** These data indicate that the reduced levels of TTSS-1 species in the ΔluxS are due to a decrease in F levels, so F is regulated by LuxS-QS mechanism. Our results agree with the experimental results presented in **[38]** Fig. 1 (A) and Fig. 2 (A).

**Δ*lsrR,* Δ*luxS* and Δ*lsrR*** [38]. When *lsrR* gene is deleted, F is expressed as in WT. However, F reaches the steady state level without delay (approximately in 3 hours) because there is no R repressing it, see S4 Fig (C) and (E). It is obvious that in Δ*lsrR* mutants, Δ*luxS* makes no difference because R represses directly F and the rest TTSS-1 species. These findings agree with the experimental results presented in [38] Fig. 2 (A).

***lsrR***$^{+}$ **plasmid** [38]. Moreover, it is confirmed by our model that overexpression of LsrR in wild-type Salmonella reduces protein levels of SPI-1. To test this case, we constructed a synthetic strain carrying a plasmid, in which expression of LsrR is not under the control of the lsr promoter (auto-repression pattern). The induction of LsrR$^{+}$ plasmid leads to overexpression of LsrR protein which in turn decreases expression of all SPI-1 species: invF, sicA, sipC were reduced by approximately 4 folds of expression in comparison with WT phenotype, see S4 Fig (C). These results confirm that LuxS-mediated quorum sensing affects regulation of invF gene, and thus virulence of TTSS-1, via LsrR regulator.

**Δ*sicA*** [36]. In sicA knock-out, we obtain WT expression of InvF protein because invF promoter is regulated by LsrR and HilA in our model. However, SipC expression is reduced about 4-folds of expression because due to the absence of SicA, there is no InvF:SicA complex formation, and thus sipC is not induced to produced SipC, S4 Fig (G). Our model confirms that in SicA knock-out there is no feedback loop in virulence circuit, as proposed in Temme et al. [36].

In order to simulate in our in silico experiments a gene knock-out, we simply have to disable corresponding reactions by nullifying their kinetic laws, see Table 1. Both in the kinetic and individual-based model, we modeled lsrFGE and lsrABCD as lsr operon (i.e. as a single unit). In an attempt to simulate knock-out of lsrFGE, we neutralized the reaction representing the $AI_2^p$ degradation by LsrF, LsrG and LsrE proteins by nullifying the pAI2_deg kinetic law (Table 1). Furthermore, to simulate knock-out of lsrB, we neutralized the reaction representing the $AI_2$ importation by LsrB transmembrane protein by nullifying the AI2_imp_ACBD kinetic law (Table 1). Finally, to simulate knock-out of luxS, in the kinetic model we introduce no $AI_2^{out}$ input pulse to the micro-environment, while in Ibm we neutralized S production reaction by nullify the kinetic law S_prod (Table 1).

In S5 Fig, we present the results for the corresponding knock-out experiment of the Ibm. We can observe that we obtain similar results to the results generated by kinetic model simulations for each lsr/TTSS-1 gene single and double knock-out experiments.

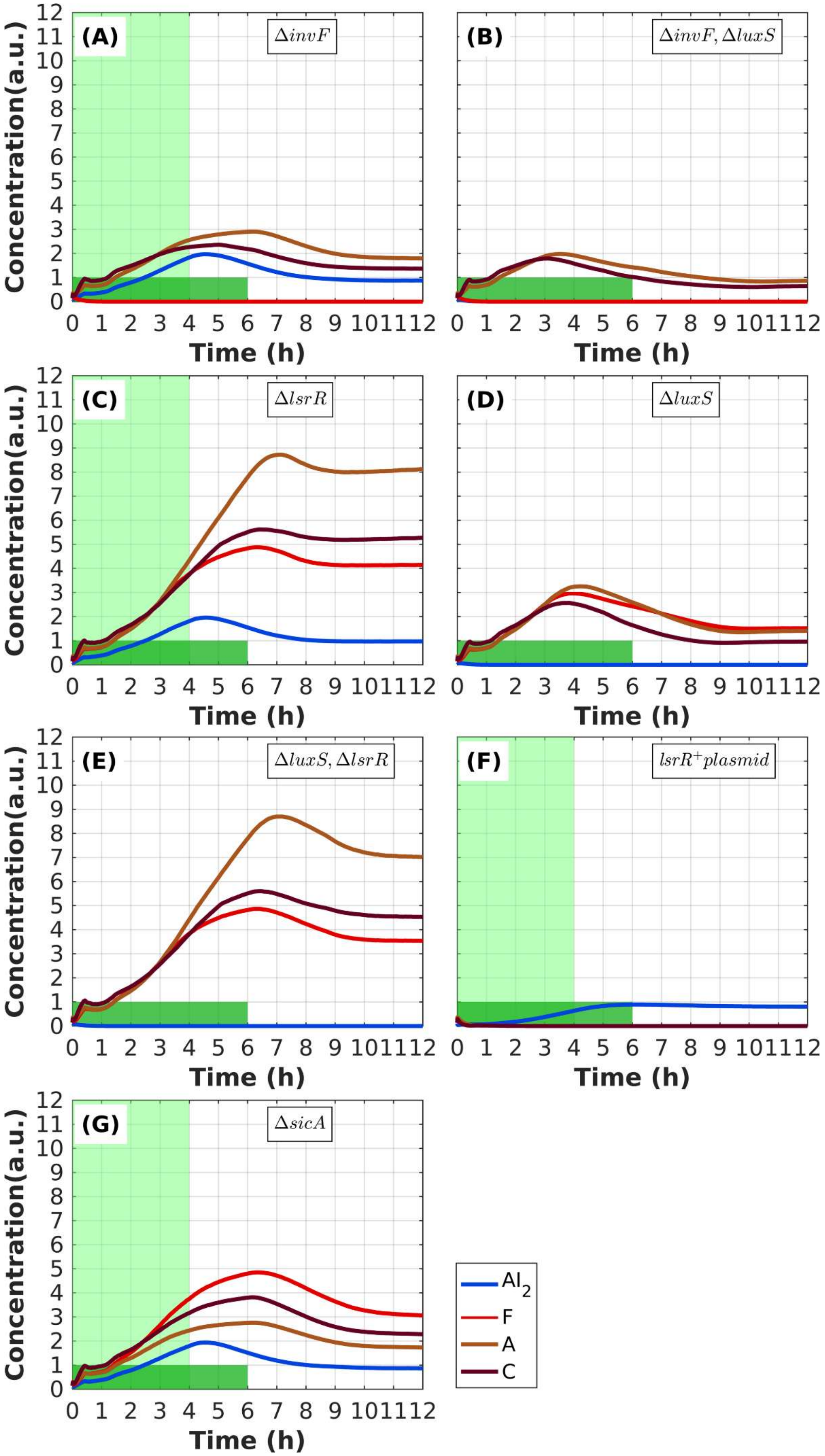

(A)
ΔinvF
(B)
ΔinvF, ΔluxS
(C)
ΔlsrR
(D)
ΔluxS
(E)
ΔluxS, ΔlsrR
(F)
lsrR+plasmid
(G)
ΔsicA
Concentration(a.u.)
Time (h)
AI2
F
A
C

**S5 Fig. lsrR, luxS, invF and sicA knock-outs effect on virulence.** Colors represent the different species. Light green region represent the interval that S is activated. Green region represent the interval that H is activated. Time-course results for 12 hours experimental time. In each knock-out in silico experiment R and TTSS-1 species fold of expression (at steady-state) matches qualitatively with the corresponding knock-out wet-lab experiments presented in *Choi et al.* [S13] and *Temme et al.* [S4]. See main text for details.

Our Ibm confirms the findings of [S4] even if it contains the QS mechanism regulating invF, extending the initial model. As we showed previously, to reproduce their finding, we varied this parameter over a physiologically relevant range, and our model confirms that the virulence persistence duration can be varied giving a remarkable diversity of relaxation times, see S6 Fig. In our experiments, we did not alter QS mechanism of WT so as to reproduce the aforementioned results.

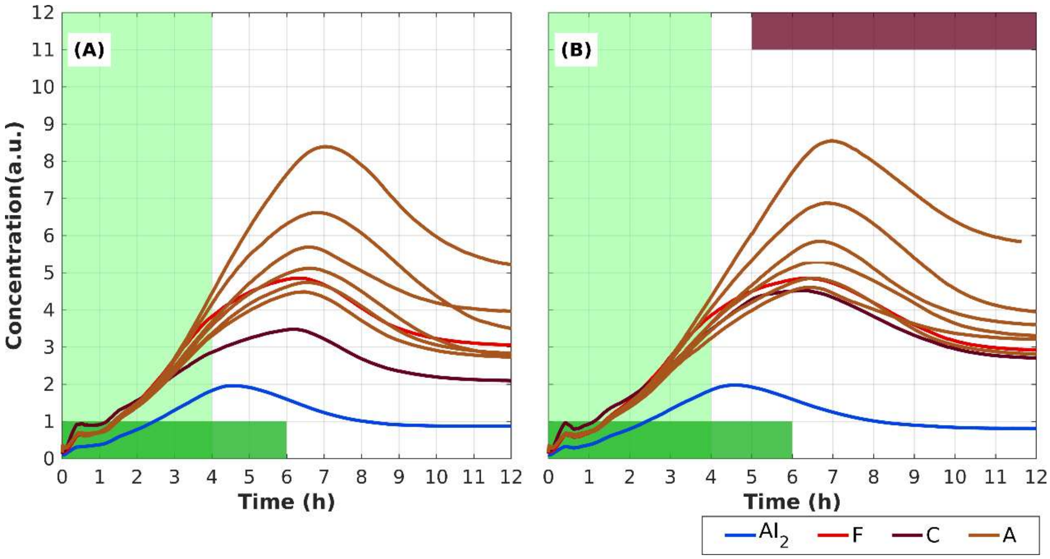


**S6 Fig. Mutations to the SicA:InvF binding site effect on A expression.** Colors represent the different species. Light green region represents the interval that S is activated. Green region represents the interval that H is activated. Marron region represent the interval that secretion is activated. Time-course results for 12 hours experimental time. Notice that when binding constant (K) increases, A expression decreases leading to lower relaxation times both in secreting and non-secreting mechanism. Here we varied K from 3.92 to 7.84 (with step 0.3922). In accordance with our kinetic model, Ibm produces similar results.

**Model predictions**

As a first testing scenario, our model is set to predict the expression of TTSS-1 species in LuxS-QS gene knock-out experiments. The results produced by the kinetic model are presented in S7 Fig.

**Δ*lsrB* / Δ*lsrB* and Δ*lsrK*.** In lsrB single knock-out and lsrB lsrK double knock-out we obtain decreasing behavior, see S7 Fig (A) and (B) correspondingly. In the former case there is very low AI-2 introduced to the system (only by PTS pathway) because there is no LsrB to import it through the transmembrane protein complex, thus TTSS-1 species expression is reduced approximately in half because LsrR repression is not fully relieved. In the latter case, TTSS-1 species expression is reduced approximately by 4 folds because there is no LsrK produced to phosphorylate AI-2, and relieve lsr operon repression by LsrR.

**Δ*lsrB* and Δ*lsrR*.** In S7 Fig (C), we did double knock-outs on lsrB and lsrR, so the TTSS-1 species are overexpressed from the beginning of the experiment, even though there is no significant concentration of AI-2 due to LsrB absence, because there is no LsrR repressing the TTSS-1 species.

**Δ*lsrFGE*.** When lsrFGE gene is knocked out, P-AI-2 is overexpressed because there are no proteins (LsrF, LsrG and LsrE) degrading it; thus, LsrR binds to it massively, loosening the repression of invF, resulting in TTSP-1 species expression as in WT (S7 Fig (D)).

**Δ*lsrK*.** In lsrK knock-out, P-AI-2 cannot be formed, thus there is no LsrR-PAI2 complex production. The LsrR represses invF. Consequently, we have reduced expression of InvF regulator, thus SicA and SipC levels are very low (S7 Fig (E)).

**Δ*lsrK*** and **Δ*lsrFGE*.** In the case of lsrK and lsrFGE double knock-out experiment, the same behavior as in single lsrK knock-out observed since there is no AI-2 phosphorylation and thus lsrFGE existence or absence makes no difference to the TTSP-1 dynamics (S7 Fig (F)).

**Δ*lsrR* and Δ*lsrK*.** In lsrR and lsrK double knock-out, we obtain early overexpression of the TTSP-1 species because LsrR cannot be produced and thus repress invF, even though there is no AI-2 phosphorylation due to LsrK absence, see S7 Fig (G).

**Δ*lsrFGE* and Δ*lsrR*.** When lsrFGE gene and lsrR genes are knocked out, TTSP-1 species are overexpressed because there is no LsrR repressing it. It is obvious that lsrFGE makes no difference in TTSP-1 species dynamics when lsrR is deleted, S7 Fig (H).

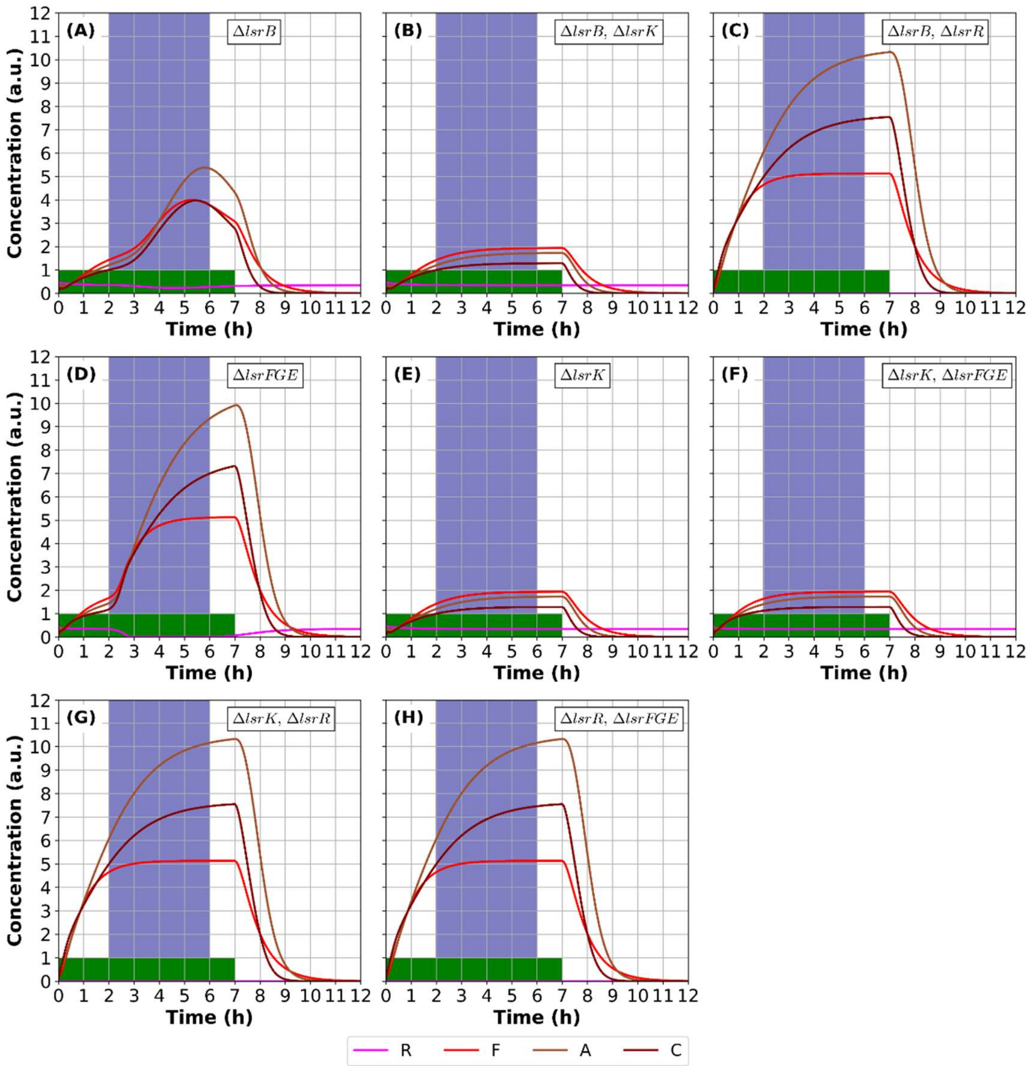


**S7 Fig. lsr genes single and double knock-outs effect on TTSS-1 species expression.** Colors represent the different species. Dark blue region represents the input $AI_2^{out}$ pulse. Green region represents the interval that H is activated. Time-course results for 12 hours experimental time. In each knock-out in silico experiment, TTSS-1 species fold of expression (at steady-state) lead to outputs that fulfill qualitatively our intuition which is based on *Choi et al.* **[38]**. However, experimental validation with the corresponding knock-out wet-lab experiments is needed. See text for details.

The corresponding results of the Ibm are provided in S8 Fig.

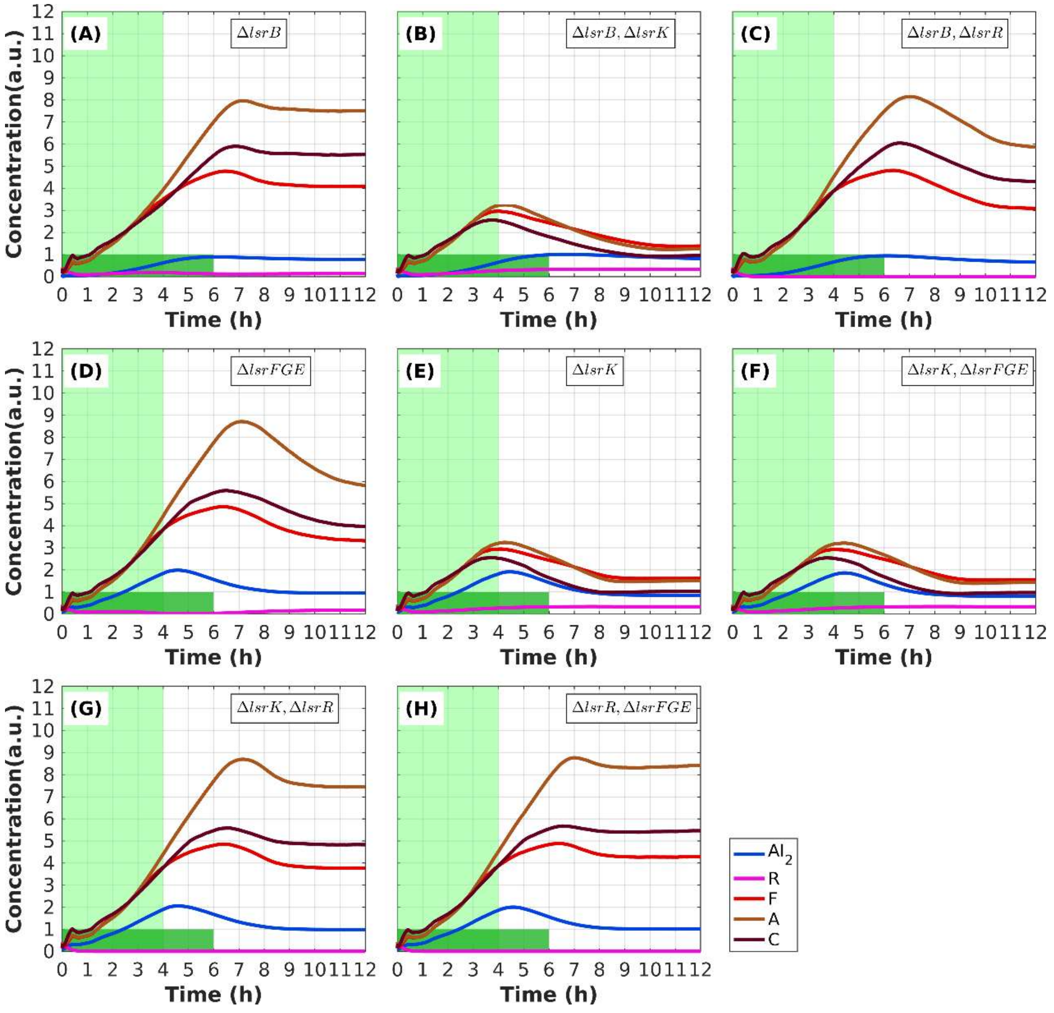


**S8 Fig. lsr genes single and double knock-outs effect on TTSS-1 species expression.** Colors represent the different species. Light green region represent the interval that S is activated. Green region represent the interval that H is activated. Time-course results for 12 hours experimental time. In each knock-out in silico experiment, TTSS-1 species fold of expression (at steady-state) lead to outputs that fulfill qualitatively our intuition which is based on *Choi et al.*[S13]. However, experimental validation with the corresponding knock-out wet-lab experiments is needed. See main text for details.

#### Nanofactories simulation with different initial cell setup

In S9 Fig, we present a simulation setup with different initial arrangement of the cells. It is obvious that given that NFs distribution, and thus $AI_2$ diffusion in the microenvironment, is the same in both simulations, the initial arrangement of the cells seems to play a more significant role in the duration that it takes to the C to be expressed. In S10 Fig (A), we present the expression of OP and C species for different number of NFs. Again, the pattern of expression increases logarithmically to the number of NFs, however in this setup the phenotype is expressed more quickly than in the aforementioned setup.

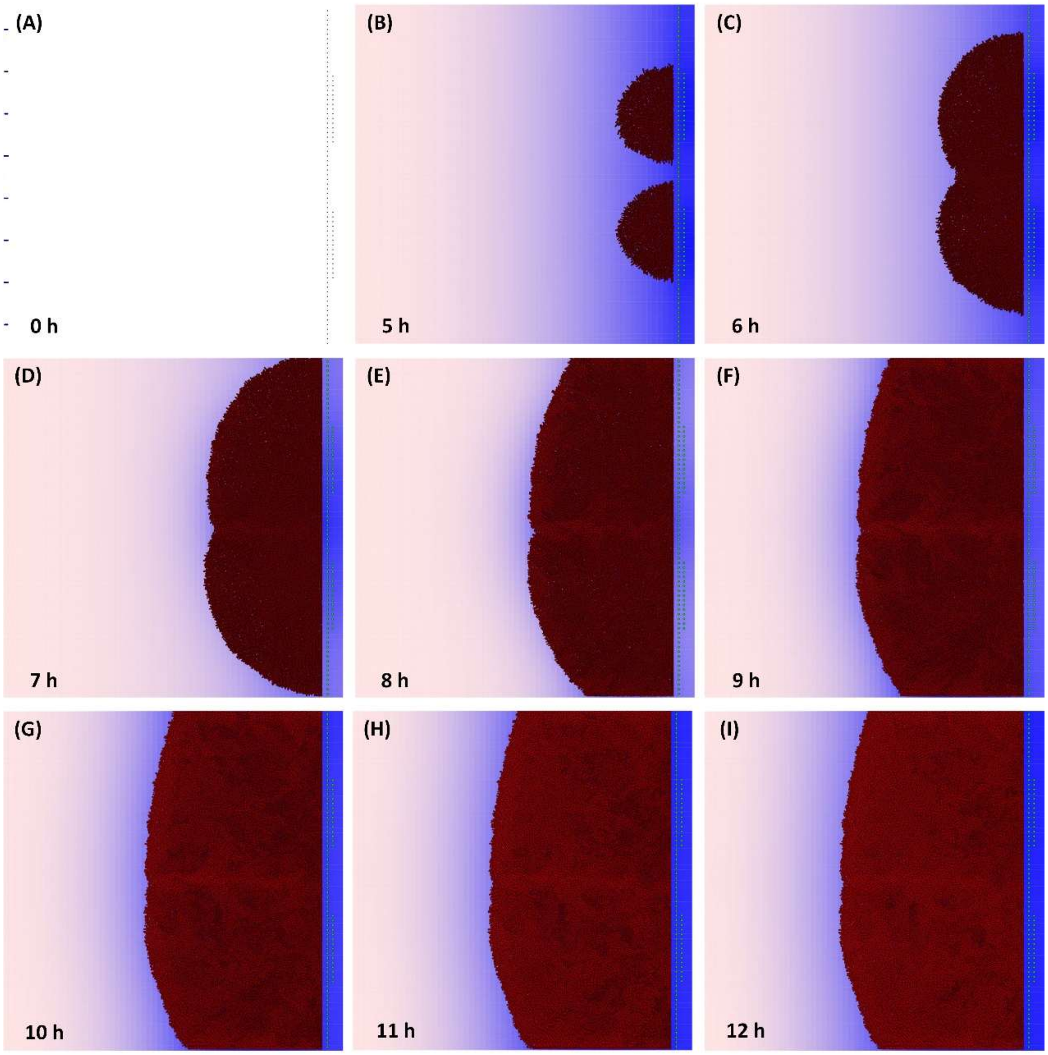


**S9 Fig. Vertical cell arrangement with uniform NFs distribution (setup 3).** (A) Initial arrangement of the simulation. At the left side of the cell surface, 8 motile cells are arranged vertically. Motile cells are able to sense $AI_2$ gradient produced by NFs. In the right side of the cell surface are the docked NFs (green circular objects). 72 NFs are distributed uniformly in the first layer while 30 NFs are distributed non-uniformly in the second layer. During the simulation, docked NFs (right side of each panel) produce $AI_2$ which in turn is diffused onto the cell surface, blue color. (B)-(I) Motile cells reach the NFs site-the source of the signal-and start to grow and express their phenotype from 5 to 12 hours. We can observe that NFs setup lead to a microenvironment with two local maxima of $AI_2$ signal. Initially the spatial pattern of the microbial community differs significantly from the spatial pattern of Setup 1. See in the S3 movie the whole simulation.

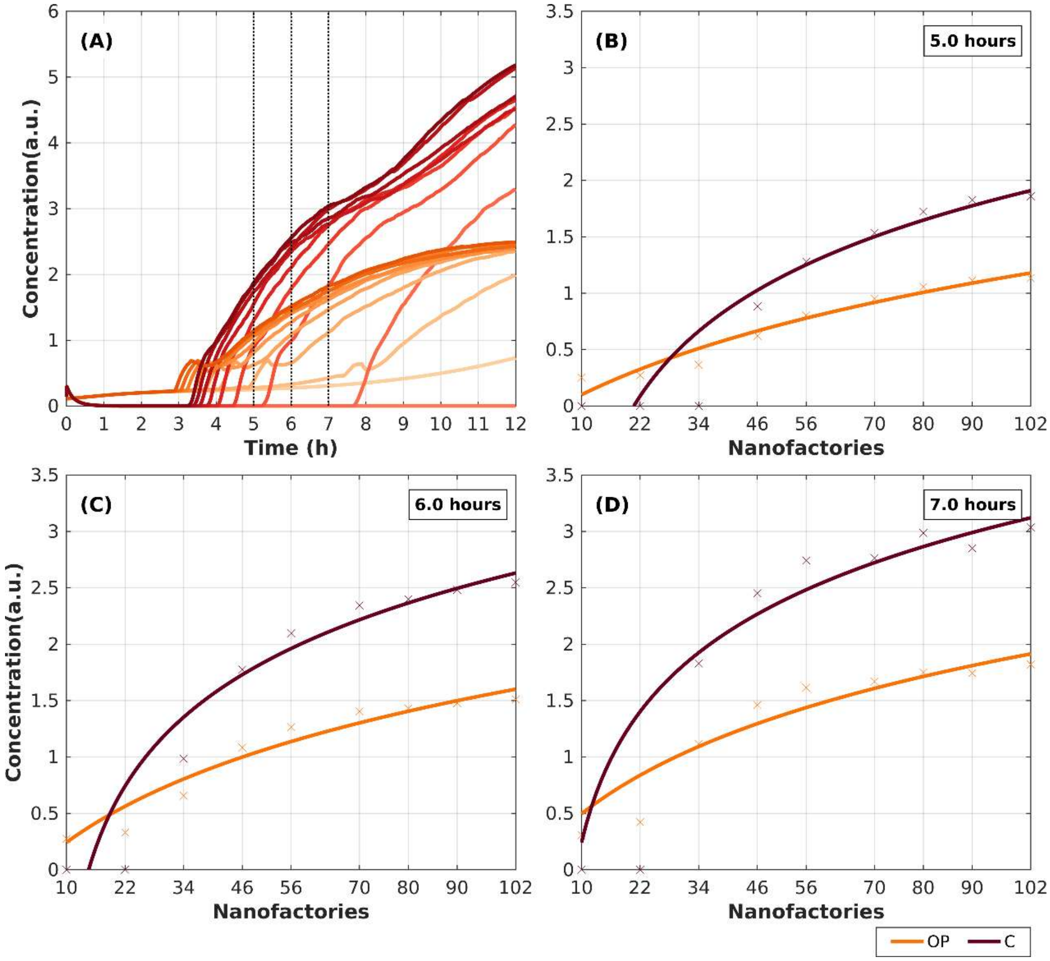


**S10 Fig. Expression of OP and C species for different number of NFs in setup 3.** Colors represent the different species. **(A)** Evolution of OP and C expression in microbial community per NF setup. Light to dark orange represent the evolution of OP from 10 to 102 NFs. Light Red to maroon represent the evolution of C from 10 to 102

NFs. (B)-(D) The expression of OP and C per NFs setup at three time points (5, 6 ad 7 hours, dashed lines in (A)). It is obvious that as the number of NFs increases the efficiency of the system increases logarithmically before the system reaches its steady state.

#### Data Availability

Due to lack of experimental data, we searched in the literature for time course experiments, concerning the species of our model. In the table below, we present the experimental data produced by different labs, found in [S1, S11, S14-S17], in similar environmental conditions on *Salmonella enterica* serovar Typhimurium strains. Specifically, all S. Typhimurium experiments concerning QS were conducted to S. Typhimurium strain ATCC 14028 and all S. Typhimurium experiments concerning T3SS SPI1 and T3SS SPI1 with QS interconnection were conducted to S. Typhimurium strain SL 14028. It is noteworthy to mention that *García-Quintanilla and Casadesús* showed that both strains are virulent, in [S18].

**S5 Table. Time course experiments.**

| Time (min) | LuxS (a.u.) | P-AI-2 (a.u.) | AI-2 (a.u.) | lsr operon (a.u.) | InvF (a.u.) | HilA (a.u.) | SicA (a.u.) | SipC (a.u.) |
|---|---|---|---|---|---|---|---|---|
| 0 | 0 | 11,9588014 | 1,1069226 | 0 | 0 | 0 | 0,00 | 0 |
| 120 | 144,024 | 48,2075325 | 2,3314855 | 7,017 | 94,96 | 51,44 | 4,53718 | 0 |
| 180 | 159,762 | 150,086355 | 151,39511 | 17,04 | 212,95 | 55,76 | 70,4421 | 241,4 |
| 210 | 157,7 | 226,24604 | 840,5649 | 24,01 | 276,2 | 57,19 | 245,802 | 345,3 |
| 240 | 154,583 | 288,25182 | 1245,9656 | 70,29 | 339,57 | 58,99 | 415,135 | 393,8211 |
| 270 | 152,9 | 126,5465215 | 740,3192 | 202 | 407,5 | 64,66 | 590,528 | 439 |
| 300 | 151,39 | 31,1789023 | 7,4351225 | 296,44 | 475,54 | 71,94 | 753,767 | 479,4922 |
| 330 | 148,9 | 20,89 | 2,811 | 266,89 | 507,9 | 77,9 | 780,40 | 521,7 |
| 360 | 147,211 | 17,5998787 | 1,5009564 | 250,26 | 540,29 | 82,73 | 789,369 | 565,1 |
| 390 | 147,1 | 10,8 | 1,5009564 | 208,71 | 540,29 | 85,47 | 782,60 | 599 |
| 420 | 147,009 | 4,024 | 1,5009564 | 204,4 | 540,29 | 88,13 | 770,459 | 612,63916 |
| **Reference** | [S14] | [S1] | [S1] | [S1] | [S15] | [S15] | [S15] | [S17] |
| **Strain** | ATCC 14028 | ATCC 14028 | ATCC 14028 SL1344 | ATCC 14028 | SL1344 | SL1344 | SL1344 | SL1344 |

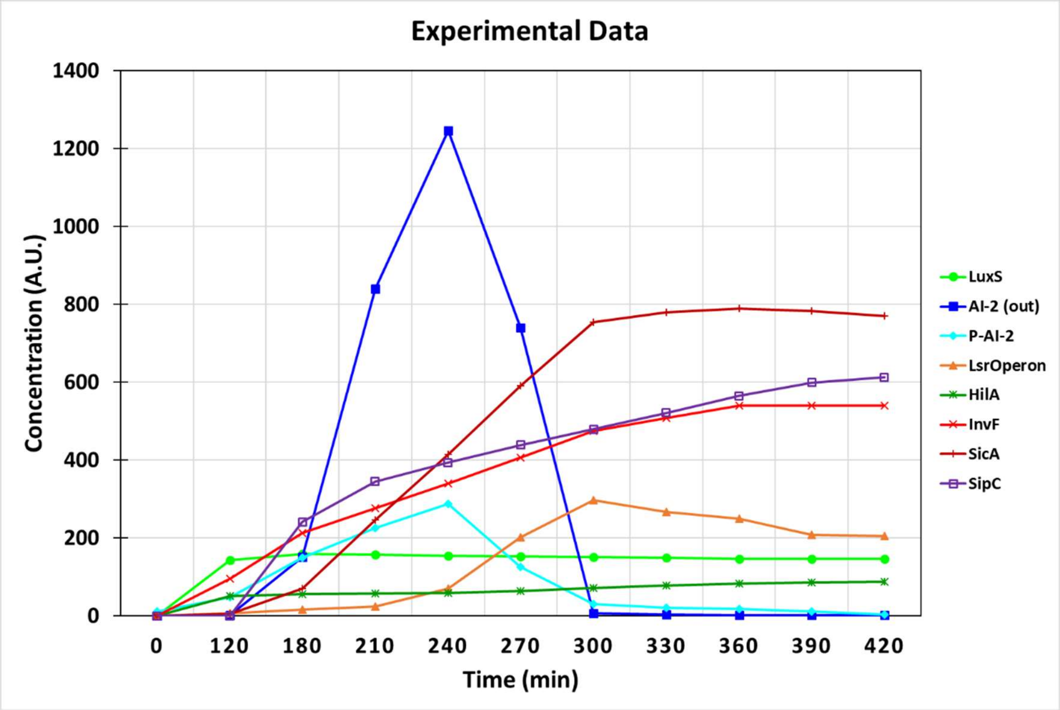


**S7 Fig. Experimental data used for parameter estimation.**

#### Parameter Estimation for kinetic model

**S6 Table. RMSE of the proposed parameter estimation approaches.**

| Algorithm | RMSE (mean parameter values) | RMSE (median parameter values) |
|---|---|---|
| Evolution Strategy | 0.864 | 1.078 |
| Particle Swarm | 0.902 | 0.937 |
| Scatter Search | 0.877 | 0.930 |
| Genetic Algorithm | 0.890 | **0.830** |

#### Parameter estimation for growth kinetics in individual-based model

Growth rate distribution parameters for each generation:

**S7 Table.**

| Generation | Alpha | Beta |
|---|---|---|
| 0 | 59.1487 | 0.0007 |
| 1 | 13.8457 | 0.0058 |
| 2 | 10.0248 | 0.0076 |
| 3 | 13.2912 | 0.0048 |
| 4 | 17.2033 | 0.0037 |
| 5 | 15.1647 | 0.004 |
| 6 | 10.0501 | 0.0038 |
| >7 | 16.4430 | 0.0025 |

Division time distribution parameters for each generation:

**S8 Table.**

| Generation | Alpha | Beta |
|---|---|---|
| 0 | 53.4126 | 2.2467 |
| 1 | 14.8704 | 2.7459 |
| 2 | 13.6637 | 2.9275 |
| 3 | 13.4675 | 3.6620 |
| 4 | 18.2327 | 2.3808 |
| 5 | 13.8790 | 3.0335 |

| | | |
|---|---|---|
| 6 | 21.8276 | 2.2974 |
| >7 | 25.2025 | 1.9167 |

Initial single-cell length distribution parameters alpha = 36.5088, beta = 0.0644.
